# Subspace based Multiple Constrained Minimum Variance (SMCMV) Beamformers

**DOI:** 10.1101/2021.05.10.443467

**Authors:** Alexander Moiseev, Anthony T. Herdman, Urs Ribary

## Abstract

In MEG and EEG brain imaging research two popular approaches are often used for spatial localization of focal task- or stimuli-related brain activations. One is a so called MUSIC approach applied in the form of RAP or TRAP MUSIC algorithms. Another one is the beamformer approach, specifically multiple constrained minimum variance (MCMV) beamformer when dealing with significantly correlated activations. Either method is using its own source localizer functions. Considering simplicity, accuracy and computational efficiency both approaches have their advantages and disadvantages. In this study we introduce a novel set of so called Subspace MCMV (or SMCMV) beamformers whose localizer functions combine MUSIC and MCMV localizers. We show that in ideal situations where forward modeling, data recording and noise measurements are error-free, SMCMV localizers allow precise identification of *n* arbitrarily correlated sources irrespective to their strength in just *n* scans of the brain volume using RAP MUSIC type algorithm. We also demonstrate by extensive computer simulations that with respect to source localization errors and the total number of identified sources SMCMV outperforms both the TRAP MUSIC and MIA MCMV (which is the most accurate MCMV algorithm to our knowledge) in non-ideal practical situations, specifically when the noise covariance cannot be estimated precisely, signal to noise ratios are small, source correlations are significant and larger numbers of sources are involved.

## 1. Introduction

A wide variety of neuroimaging studies using magnetoencephalography (MEG) and electroencephalography (EEG) involve identifying a set of focal, spatially compact task or stimuli-related activations inside the brain volume. Achieving this by means of an array of sensors positioned outside the brain is referred to as a bioelectromagnetic inverse problem [1]. Very often time courses of these activations are correlated with each other because different parts of the human brain are working together performing a task or responding to external events. In some situations the correlations become very strong, such as in the case of stimuli-driven activation of bilateral sensory areas (see for example [2, 3]). Typically existence of strong time course correlations make solution of the inverse problem much harder.

When the number of activations, or brain sources of interest is relatively small compared to the total number of sensors in the M/EEG array (so called “low rank” activity), two popular techniques are often applied. Both have their advantages and drawbacks.

The first one called Multiple Signal Classification (MUSIC) [4] is based on a separation of a linear vector space spanned by the recorded data (also called the “sensor space”) into two orthogonal subspaces: a source, or signal subspace, and a noise subspace. Strictly speaking this is possible when the noise background is spatially white; in a general case pre-whitening of the data is necessary. Assuming that the sources of interest are current dipoles, the problem of identifying them is then solved by testing if a lead field of a source in a probe location belongs to the source subspace. A commonly used practical way to do this is called a Recursively Applied and Projected (RAP) MUSIC [5, 6], where sources are found by iteratively searching the brain volume for maxima of a specially designed function.

Such functions whose extrema (by convention - maxima) identify the sources are called localizer functions, or just “localizers”. They are constructed based on the bio-electromagnetic brain model and the observed data. A localizer is unbiased if its maxima match the source parameters precisely. The RAP MUSIC localizer is unbiased, as shown in *Appendix A*.

A significant improvement to the RAP MUSIC procedure was recently suggested in [7], which addresses a problem of additional spurious sources often reported by the original algorithm [5, 6]. Another important note is that the MUSIC approach by itself only locates sources but does not provide their time courses. However reconstructing those is a much easier task and can be done as in the second technique described below.

This another technique known as minimum variance beamforming originates in idea of constructing a “spatial filter”, that is a linear combination of M/EEG sensor channels which amplifies brain activity from location of interest and at the same time suppresses interfering signals from other places. It turns out that coefficients of such a combination (i.e. filter weights) can be found in a simple form (see [8, 9, 10, 11] and Eq.(2) below). Moreover this spatial filter yields the best time course reconstruction in a mean squared error sense provided that the probe location and orientation match those of the true source [11]. When *a priori* information about source positions and orientations is missing, the brain volume, or more generally the task parameter space is searched for sources using localizer functions similarly to the MUSIC case. For beamformers the simplest expressions for the localizers can be obtained assuming that only a single active source plus uncorrelated noise exist [11]. However in practice the same expressions are often applied in situations with multiple brain activations. This constitutes a traditional *single source* beamformer approach.

In case of multiple simultaneous activations single source localizers are biased. Even worse, when the source time courses are strongly correlated, localizer maxima may be suppressed or disappear completely, because with respect to each source an assumption of uncorrelated noise interference is violated. This is usually called the correlated sources cancellation problem [9, 12, 13]. The latter can be addressed by multi-source beamformer approaches where all sources are considered simultaneously. Specifically, in multiple constrained minimum variance (MCMV) beamformers [8, 14, 15, 16, 17] each source weight is calculated subject to constraints that filter gains for signals originating from other source locations, or even from extended brain regions are zero. This would solve the correlated interference problem for time course reconstructions if locations and orientations of all sources could be identified, but as mentioned above single source localizers cannot be used for that purpose. To address this a set of unbiased *multi-source* MCMV localizer functions was derived in [17] (also see section 2.3 below). In contrast to the single source approximation those localizers depend on positions and orientations of all sources at once therefore searching for extrema occurs in a parameter space of many dimensions. A “brute force” exhaustive search which was computationally affordable in a single-source case is no longer an option. For this reason approximate iterative source search procedures were suggested [17, 18].

Formally a big advantage of the MUSIC approach over the minimum variance beamformers is that no assumptions about source correlations are necessary, thus one could expect it to work equally well for both correlated and uncorrelated sources. Moreover in an ideal situation (T)RAP MUSIC should yield precise source locations in just *n* consecutive scans of the brain volume, where *n* is the number of sources. A disadvantage of this approach is that its success solely depends on accurate separation of the sensor space into the signal and noise subspaces. Once this is achieved the source is identified simply by the subspace containing its lead field – in other words localization becomes purely geometrical. All other factors such as source magnitude or signal to noise ratio (SNR) are ignored. Actually the TRAP improvement [7] is a way to deal with this issue. In practice precise separation is hard to achieve, even more so with strong source correlations – thus in fact the MUSIC approach is not immune to the source correlations problem.

In contrast, both traditional and multi-source beamformer localizers seek to achieve maximal SNRs of the reconstructed sources. Note that in many situations getting the best SNR is the most important criterion of the quality of the reconstruction. Moreover MCMV beamformers can deal with strong source correlations. However while theoretically unbiased, due to high dimensionality of the parameter space MCMV beamformers require approximate iterative search procedures [17], the most accurate ones being rather complicated and time consuming [18] and still not necessarily converging to the true solution.

In this work we introduce a generic family of novel multi-source localizer functions combining both the above approaches and addressing the disadvantages of either technique. They allow establishing precise locations and orientations of a (low rank) set of *n* arbitrarily correlated brain sources irrespective to their SNRs in the absence of forward modeling, data and noise covariance estimation errors – in other words, they are unbiased. Localization is achieved in just *n* searches over the brain volume. In non-ideal practical situations source localization is still robust because SNR is included in the target cost functions, which makes errors in separating the sensor space into signal and noise subspaces much less damaging. Our theoretical results are validated and compared to both MUSIC and MCMV approaches by means of extensive computer simulations based on real EEG and MEG data.

## 2. Methods

We start with a formulation of the M/EEG inverse problem and give a brief overview of RAP/TRAP MUSIC and MCMV approaches. We then proceed with introduction of the subspace based (SMCMV) localizers and discussion of their properties.

Throughout this paper, the following notation is used. Vectors and matrices are defined with bold lower and uppercase letters, respectively, while scalar quantities - with regular letters. Angle brackets denote statistical averaging; expression 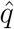 refers to an estimate of some quantity, as opposed to its true value *q*; 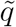 denotes the fluctuating part of *q*: 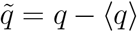. Superscript “*T*” means matrix transposition; the “dagger” symbol “†” denotes matrix pseudo-inverse. Generally we follow the notation adopted in [17, 18] as some of the results are used in this study.

### 2.1. M/EEG measurements model

We assume that M/EEG signals are recorded by an array of *M* sensors. Let ***b***(*t*) denote a *M*-dimensional column vector of sensor readings at time *t*, and suppose that ***b***(*t*) is a sum of signals produced by *n << M* sources of interest with time courses *s*(***θ**^i^, t*)*, i* = 1*, …, n*, and noise ***ν***(*t*). Here ***θ**^i^* denotes a set of parameters defining the source *i*. For dipole sources ***θ**^i^* = {***r**^i^, **o**^i^*}, where ***r**^i^* is the source location in 3D space and ***o**^i^,* ∥***o**^i^*∥ = 1 – its orientation. Then combining all source time courses in a *n*-dimensional vector ***s***(**Θ***, t*) = {*s*(***θ**^i^, t*)}^*T*^, **Θ**= {***θ**^i^*}*, i* = 1*, …, n*, and adopting standard assumptions for the bio-electromagnetic forward problem [1], we can write

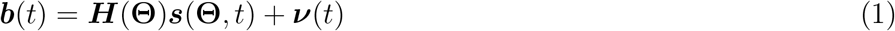

In Eq.(1), (*M* × *n*) matrix ***H*** = {***h***(***θ**^i^*)}*, i* = 1*, …, n* is comprised of the *M*-dimensional column vectors ***h**^i^* = ***h***(***θ**^i^*) of the true source lead fields. We assume that functions ***h***(***θ***) are known, ***θ**^i^* do not change with time, (***ν***(*t*)) = 0 and that processes 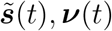 are stationary and uncorrelated. Note that in this work we do not presume any *a priori* dependence of the source orientations on locations ***r**^i^* (so called “freely oriented dipoles”). By definition ***ν*** describes that part of the measured field ***b*** which is not produced by the sources of interest and thus includes the environmental and instrumental noise as well as the fields generated by all other brain sources (the brain noise). For the purposes of this study the only information we need about the noise field is its (*M* × *M*) covariance matrix ***N*** = (***νν**^T^*). An accurate estimate of ***N*** is crucial for all the approaches discussed below, and in practice constitutes one of the biggest challenges. The number of sources of interest *n* depends on the situation and may or may not be known in advance. In the latter case we assume that *a reasonable upper boundary for n is known a priori* and this number is being used in calculations.

Reconstructing the brain sources from the measurements ***b***(*t*) is referred to as the bio-electromagnetic inverse problem [1], [11], [19]. It may be solved by first localizing the sources, that is by finding the parameters ***θ**^i^*, then by estimating corresponding time courses *s*(***θ**^i^, t*). In this study we only focus on the localization step, because once ***θ**^i^* are established, the best possible estimate of the time courses ***s***(**Θ***, t*) in the root mean square (RMS) sense is well known. This is the minimum variance beamformer expression [8, 11]

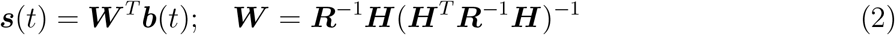

In Eq.(2) ***W*** = {***w**^i^*}*, i* = 1*, …, n* is a (*M* × *n*) matrix of the beamformer weight vectors ***w**^i^*, and 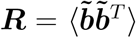 is the (*M* × *M*) data covariance matrix; we dropped the dependence of ***s**, **H*** and ***W*** on **Θ** for brevity.

In the majority of practical situations the main contribution to the noise field ***ν*** comes from sources in the brain itself and other signals originating in the subject’s head, such as cardio and muscle artifacts and eye blinks. Those signals have complex spatial structures. This results in covariance matrix ***N*** being quite different from the diagonal white noise covariance ***N**_white_* = *σ*^2^***I**_M_*, where *σ* is the RMS noise level in a single sensor channel and ***I**_k_* denotes a *k*-dimensional identity matrix. While for MCMV solutions discussed below specific form of ***N*** is not important, the MUSIC-based approaches strictly speaking are only valid under the assumption of the diagonal white noise. To use those in case of an arbitrary ***N*** a pre-whitening transformation is required. Specifically, the following change of variables is performed:

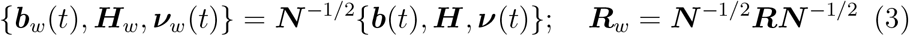

The model equation in the “pre-whitened” variables ***b**_w_*(*t*)*, **H**_w_, **ν**_w_*(*t*) has exactly the same form (1) but now the noise covariance becomes an identity matrix: 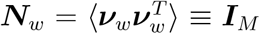. *From here on we assume that transformation (3) is applied and the pre-whitened variables are used everywhere*, omitting subscript “*w*” for brevity. However we do not substitute the identity matrix and keep using symbol ***N*** for the noise covariance in cases where expressions hold in a general non-diagonal case.

### 2.2. RAP and TRAP MUSIC

The MUSIC methods [4, 6, 7] are based on the fact that in case of diagonal white noise the *M*-dimensional sensor space spanned by vector ***b*** can be split into two orthogonal subspaces: the *signal* or *source* subspace, and the *noise* subspace. The signal subspace is defined as a column space of (*M* ×*n*) matrix ***U*** = {***u**^i^*} comprised of the first *n* eigenvectors of the (pre-whitened) data covariance matrix ***R***: ***Ru**^i^* = *σ_i_**u**^i^,* ||***u**^i^*|| = 1*, i* = 1*, …, n* where eigenvalues *σ_i_* are sorted in a descending order; it can be shown that *σ_i_ >* 1. The remaining *M* −*n* eigenvectors span the noise subspace and all have eigenvalues equal to 1: *σ_i_* = 1*, i* = *n* + 1*, …, M*. Importantly, all the source lead fields ***h**^i^* strictly belong to the source subspace and are orthogonal to the noise subspace. This allows to construct so called *MUSIC localizer* function as follows.

For any brain location ***r***, define (*M* ×3) “vector” lead field matrix ***L***(***r***) = {***h**^x^, **h**^y^, **h**^z^*} where ***h**^x,y,z^* are lead fields of point dipoles at ***r*** oriented along *x, y, z* axes of the Cartesian coordinate system. Then the lead field of a dipole with an arbitrary orientation ***o*** is ***h***(***r**, **o***) = ***L***(***r***)***o***. Let ***P*** be a projection operator on the signal subspace: ***P*** = ***UU** ^T^*, then for any vector ***v*** in the sensor space its projection on the signal subspace is ***P v***. In the freely oriented dipoles case, the MUSIC localizer *μ* is defined as the largest eigenvalue of a 3D generalized eigenvalue problem:

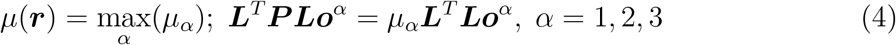

Physically *μ*(***r***) represents a maximum value of a squared ratio of the length of a projection ||***P h***(***r**, **o***)|| to ||***h***(***r**, **o***)|| – that is, of the square of cosine of the angle between ***h*** and the signal subspace – over the source orientations ***o*** at the probe location ***r***. Three-dimensional unit eigenvector ***o**^α^* represents the orientation where the maximum is reached. *μ*(***r***) equals 1 for the true sources because their lead fields belong to the signal subspace, and is less than 1 for other brain locations. Thus the sources of interest can be found by scanning the brain for maxima of *μ*(***r***); moreover those maxima point to the true source locations for an arbitrary signal to noise ratio (SNR) and arbitrary inter-source correlations.

In practice due to measurement and modeling errors a more reliable way is to localize the sources iteratively. The RAP MUSIC algorithm [6] starts with scanning the brain for the largest value of *μ*; corresponding 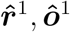 are assumed to be the parameters of the first source. At the *k*-th iteration, *k* − 1 sources are already localized, and the lead field estimates 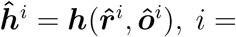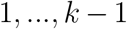 are available. Define (*M* × (*k* − 1)) matrix 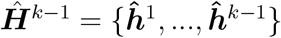 and construct an operator

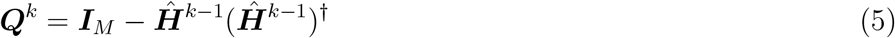

which projects any vector in the sensor space on a subspace orthogonal to all 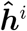. Consider (*M* × *n*) matrix ***S**^k^* = ***Q**^k^**U***. Its span represents the original signal subspace from which the first *k* − 1 sources were (approximately) out-projected. Define a projection operator on this new subspace: ***P** ^k^* = ***S**^k^**S**^k^*^†^. Then the *k*-th source in the RAP procedure is found by scanning the brain with the same localizer (4) where the original projector ***P*** is replaced with ***P** ^k^* and the lead field ***L*** is replaced with ***Q**^k^**L***:

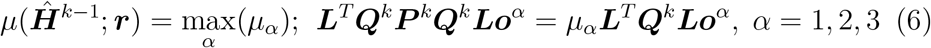

An important improvement to the RAP MUSIC algorithm was recently introduced in [7]. Note that if the operator ***Q**^k^* were constructed using the true source lead fields {***h***^1^*, …, **h**^k^*^−1^}, the “out-projected” source subspace ***S**^k^* would have *n* − *k* + 1 dimensions. In practice the estimates 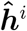 are never exactly equal to the true ***h**^i^*. Consequently the residuals of the true sources ***Q**^k^**h***^1^*, …, **Q**^k^**h**^k^*^−1^ are small but not zero and belong to ***S**^k^*, thus the latter subspace still has *n* dimensions. Importantly, these small residuals produce strong localizer maxima in the vicinity of the already found sources, because from (6) it follows that 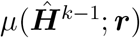 only depends on the angle between the probe (in this case – the residual) lead field ***Q**^k^**h*** = ***Q**^k^**Lo*** and the source subspace, and ignores its magnitude [7]. As a result, RAP procedure tends to find additional false sources. A solution to this problem suggested in [7] and called “Truncated RAP” or “TRAP” MUSIC is to manually remove *k* − 1 residual dimensions of ***S**^k^* before searching for the *k*-th source. This is achieved by performing a singular value decomposition of matrix ***S**^k^* and setting its *k* − 1 smallest singular values to 0. The resulting “truncated” subspace 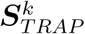 is used to construct the projector 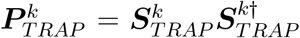, which is substituted into Eq.(4) instead of ***P***.

### 2.3. MCMV and RAP beamformers

Inverse solutions referred to as minimum variance beamformers seek the source time courses in the spatial filter form given by the 1^st^ equation in (2). The weights themselves (the 2^nd^ equation) are obtained by minimizing reconstructed signal power under certain constraints (hence the name “minimum variance beamformers”). The original formulations [8, 9, 10, 11] applied this principle to each source separately by writing *s*(***θ**^i^, t*) = ***w**^T^* (***θ**^i^*)***b***(*t*) and getting for the weights ***w***(***θ**^i^*) = ***R***^−1^***h**^i^*(***h**^iT^ **R***^−1^***h**^i^*)^−1^; *i* = 1*, …, n*. This differs with solution (2) in that weights ***w**^i^* = ***w***(***θ**^i^*) for each source depend solely on its own parameters ***θ**^i^* ignoring existence of all other sources – this is why it is conventionally called a *single source* beamformer. The traditional single source beamformers are widely used by neuroimaging community and in many cases provide excellent results. However the single source approach has important limitations. First, maxima of corresponding localizer functions do not point exactly to the source locations unless *n* = 1 so the localizers are biased for *n >* 1. Second, as mentioned in the *Introduction*, when *n >* 1 and source time courses are correlated, those maxima become smaller or may disappear (the “correlated sources cancellation” problem). Third, strong source correlations result in distorted source time courses.

These problems are resolved by a Multiple Constrained Minimum Variance (MCMV) beamformer, which in various forms was suggested, for example, in [14, 15, 3, 16]. In all cases the filter weights ***W*** = {***w**^i^*} are given by Eq.(2). Their important property is that each weight ***w**^i^* is orthogonal to lead fields of all other sources ***h**^j^*: ***w**^iT^ **h**^j^* = 0*, j* ≠ *i*, or in matrix form: ***W** ^T^ **H*** = ***I**_n_*. Physically it means that individual spatial filters ***w**^i^* have nulls in the directions of other sources, which prevents source cancellation and time courses corruption.

To apply the MCMV filter (2) in practice one needs to know all sources parameters **Θ** but as explained above one cannot use single source localizers for this purpose. This problem was addressed in [17] where unbiased multisource localizer functions were found. Three of those used in this work are defined as follows [17]:

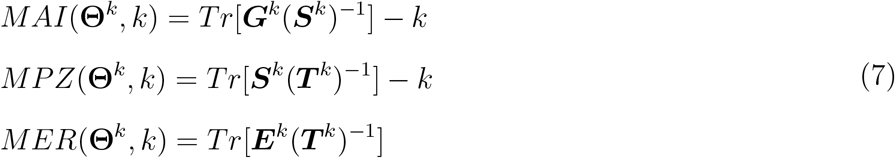

The abbreviations above stand for Multi-source Activity Index (MAI), Multisource Pseudo-Z (MPZ) and Multi-source Evoked Response (MER) localizer, respectively. In Eq.(7), for any set of *k* probe sources (*k* × *k*) matrices ***G**^k^, **S**^k^, **T** ^k^, **E**^k^* are given by equations

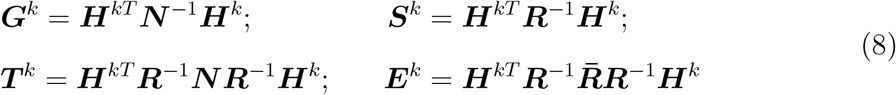

and ***H**^k^* is the (*M* × *k*) probe lead field: ***H**^k^* = {***h***^1^*, …, **h**^k^*}. The (*M* × *M*) matrix 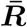 contains 2nd moments of the epoch-averaged (i.e. *evoked*) fields 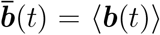, time-averaged over a certain interval of interest: 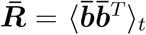. Equations (7), (8) contain ***N*** explicitly because they are valid in both the original and pre-whitened coordinates. For *k* = 1 they are reduced (up to an additive constant) to known expressions for the single source scalar beamformers.

As proven in [17], localizers (7) achieve a global maximum over a parameter space {**Θ***^k^, k*} when *k* becomes equal to the true number of sources *n* and all ***θ**^i^, i* = 1*, …, n* match the parameters of the true sources. Once the maximum is reached, making *k > n* does not increase the localizer value any more.

A brute force search for extrema of expressions (7) is computationally prohibitive unless the number of sources is just one or two. Therefore approximate iteration procedures were suggested in [17, 18] which replace a search in the high-dimensional parameter space with a number of consecutive searches of the conventional 3D brain space.

Specifically, in a Single-step Iterative Approach (SIA) one starts by setting *k* = 1 and finds parameters of the 1st source 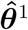 as the largest maximum of corresponding localizer (7) over the brain volume just as in the single source case. At the *k*-th iteration step estimates 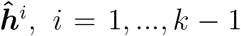 are available. To find the *k*-th source construct (*M* × *k*) lead field matrix 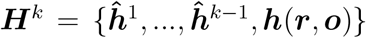, where ***h*** is the probe lead field. ***H**^k^* is then plugged into (7), and the estimates 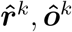 of the next source location and orientation are determined from the largest maximum of corresponding MCMV localizer by varying only ***r*** and keeping all 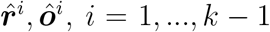 fixed. For each ***r*** the optimal orientation ***o***(***r***) is defined by expressions found in [17].

An alternative iterative procedure called a *RAP beamformer* was suggested in [19]. Similar to SIA, the RAP beamformer finds the first source parameters 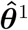 using a single-source localizer. At *k*-th step an out-projector ***Q**^k^* of already found *k* − 1 source lead fields is constructed as in Eq.(5) and applied to the forward equations (1). The *k*-th source is again found using the single source localizers which are now applied to the reduced rank out-projected data. RAP beamformer expressions corresponding to MAI and MPZ were derived in [19]; the RAP version of the evoked localizer MER is obtained identically (cf. (7)):

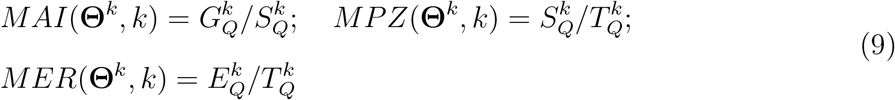

where 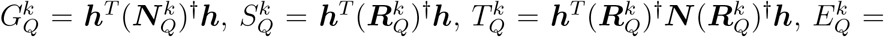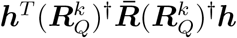 and 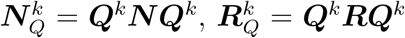. Maxima of (9) are found by varying the probe location ***r***; orientation ***o***(***r***) is determined by solving corresponding 3D eigenvalue problem as is well known for the single source beamformers [11, 19].

SIA and RAP beamformer algorithms are not equivalent but closely related, and typically both result in similar series of the estimated source parameters 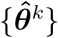*, k* = 1*, …, n* [19]. Both require exactly *n* scans of the brain volume to extract *n* sources and corresponding computational effort is approximately the same as for the RAP/TRAP MUSIC procedure.

In contrast to SIA and RAP beamformers, the Multi-step Iterative Approach (MIA) suggested in [18] does not fix already found sources when searching for maxima of localizers (7), and uses additional iteration loops to update their parameters after each new candidate source is added. As shown in [18], MIA significantly outperforms SIA in accuracy but is much more computationally intensive.

### 2.4. SMCMV localizers

In theory both MUSIC and MCMV localizers allow finding all sources precisely in error-free situations for arbitrary SNRs and inter-source correlations. However either have their advantages and disadvantages. An advantage of the RAP/TRAP MUSIC approach is this potentially precise reconstruction is achieved using a simple single loop iteration procedure. In contrast SIA or RAP single loop beamformer solutions yield approximate results even in an ideal case. On the other hand disadvantages of the MUSIC localizer are that it requires precise signal subspace identification and is purely “geometrical”, thus prone to detecting spurious sources (see the discussion following Eq.(6) above). At the same time MCMV localizers are based on SNR so candidate sources with low SNRs are penalized by construction. As a result in practice even single source beamformers often turn out to be more accurate than the MUSIC one.

In this section we introduce novel localizer functions called *”Subspace MCMV”* (SMCMV) localizers, which combine the advantages of both the RAP MUSIC and the MCMV approaches. They are based on an observation that any function of a single probe source parameters ***θ*** of the form *f* (***θ***) = (1 − *μ*(***θ***))*P* (***θ***), where *μ* is the MUSIC localizer (4) and *P* – some smooth positive function, achieves a local minimum when ***h***(***θ***) matches a true source. Thus we can choose *P* (***θ***) so that *f* (***θ***) becomes an unbiased localizer weighted by source parameters of our choice, such as source SNR. We also want an efficient procedure for extracting all the sources in a predictable number of iterations, and conventionally to have localizer maxima rather than minima to identify the sources.

These considerations lead to the following generic definition of an iterative SMCMV localizer. At each iteration step 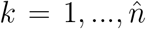 let 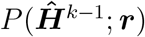 be a smooth limited positive function of a set of *k* − 1 already estimated source lead fields 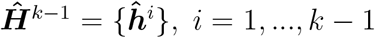 in pre-whitened coordinates, and a probe brain location ***r***. Then SMCMV localizer is defined by expression

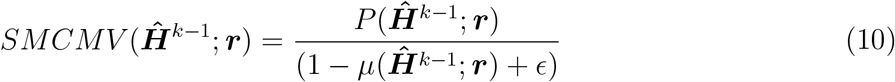

where 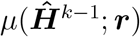 is the RAP MUSIC localizer (6) and *ϵ* is an arbitrarily small positive number. The latter is only needed to avoid a singularity in an ideal case of precise localization where *μ* = 1. In practice it can be omitted because *μ* never reaches 1. The brain location corresponding to the maximum of (10) is taken as the next source position 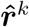; orientation 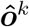 is defined by expression (6).

Specific versions of SMCMV localizers are obtained by substituting for 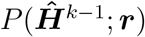 single loop iterative solutions for MAI, MPZ and MER beamformers, such as SIA versions of MCMV (7), (8) or RAP expressions (9). Other possible choices are briefly touched in *Discussion*.

It is clear from Eq.(10) that SMCMV can be viewed as the RAP MUSIC solution modulated by MCMV localizer, that is by the signal-to-noise ratio.

Equivalently it can be regarded as a MCMV single loop iterative solution with a “geometrical” correction given by the MUSIC localizer.

Importantly, *any SMCMV localizer (10) provides unbiased estimate of the source parameters for arbitrary SNRs and inter-source correlations*, as shown in *Appendix A*. By construction this only requires 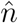 3D searches for a global localizer maximum over the brain space. Thus SMCMV computational cost is similar to that of RAP/TRAP MUSIC, SIA MCMV or RAP beamformer procedures. At the same time unlike the former in the SMCMV case the source SNRs are taken into account. Also in contrast to the SIA MCMV or RAP beamformer the iterations yield exact results in error-free situations. We investigate SMCMV performance in the following sections.

## 3. Computer simulations: setup and analyses

Since SMCMV combines MUSIC and MCMV approaches, we wanted to compare its performance to benchmark MUSIC and MCMV solutions. We chose TRAP MUSIC [7] to represent the first category because it provides significant improvements compared to the traditional RAP MUSIC. We chose MCMV MIA algorithm [18] for the second because to our knowledge this is the most accurate MCMV localization procedure available by now.

Reliable performance estimates can only be done when the ground truth is known, therefore numerical modeling is the only option. One also needs realistic settings where errors are present. To be as close to practical situations as possible we used real human head models, M/EEG data recorded in real experiments to represent the brain noise, estimated data and noise covariance matrices over limited time intervals, and introduced modeling errors as described in the following sections.

### 3.1. Simulations setup

Both MEG and EEG modalities were tested.

In the MEG case simulated datasets were generated for a 151-channel axial gradiometer CTF MEG system (Canada). The head model for an adult human subject was computed by extracting a brain hull surface from subject’s MRI data; multiple local spheres method [20] was used to generate current dipole forward solutions. 10 minutes of continuous resting state MEG data were collected for this subject at 200 samples per second with third gradient environmental noise cancellation in [0.5 - 50] Hz frequency range; these data were used as the noise background.

The source activity was added to the noise by planting randomly oriented current dipole sources in randomly chosen voxels of a rectangular grid with a 2 mm step (the *”source grid”*), and projecting their fields to the sensors. Each source time course consisted of 300 one second long narrow band pulses with one second intervals between them. Pulses had 20 Hz central frequency, 2 Hz bandwidth, Hanning envelopes, and were triggered synchronously for all sources. Thus we had 300 two seconds long epochs modeling the source activity. For every epoch individual source waveforms were tuned to obtain desired inter-source correlation matrix. To model the evoked (phase locked) activity all epochs had identical source waveforms; in case of the time-locked activity a different set of pulses with the same inter-source correlations was generated for every epoch.

MEG inverse solutions were obtained by calculating localizer values for each voxel of a rectangular grid with a 5 mm step (the *”measurement grid”*). The voxels of the source and measurement grids did not coincide so the true sources could be arbitrarily positioned relative to the measurement grid voxels, introducing the forward modeling errors.

For the EEG case we used publicly available data recorded by a Neuroscan EEG system with 55 working channels [21]. Specifically, 226 4.095 seconds long epochs of EEG data collected at 200 Hz sampling rate were pre-processed by applying a high pass filter with 2.5 Hz cut off. To be used as a brain noise background the data was additionally randomized to ensure that no effects of task related activity were present while preserving frequency power spectra of all sensor channels and the spatial covariance of the data, as explained in *Appendix B*.

The simulated sources were added to this noise background the same way as for the MEG data but now intervals between pulses were 3.095 s due to a longer epoch length. The source grid with 3 mm step was used to position the sources for forward modeling, while 5 mm step measurement grid not matching the source grid was used for reconstructions. In both cases the lead fields were generated for Colin MRI template using a three layer BEM model with standard conductivity values as implemented in Brainstorm and OpenMEEG software [22, 23].

We evaluated performance of all the methods depending on the modality (MEG/EEG), the number of active sources, their SNRs observed at the sensor array, inter-source correlations, and quality of the noise model used. The performance parameters we looked at were average spatial localization errors and average numbers of lost (i.e. not found) sources for each technique. The source was considered lost if no candidate sources were found within a 2 cm radius from the known true source position.

The number of sources tested were *n* = 2, 5, 10 and 20 for MEG; for EEG the 20 sources setting was omitted because of violation of condition *n << M*. In all cases the *a priori* hypothesized number of sources 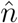 exceeded the true *n* by 2. We found that the results were almost insensitive to 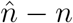 if it was within reasonable limits (2 - 5).

The sensor level *amplitude* SNR for a source with a lead field ***h*** was defined by formula *SNR* = ⟨*s*^2^⟩^1*/*2^||***h***||*/T r*(***N***)^1*/*2^. The following SNR values were tested for MEG: 0.05, 0.1, 0.2, 0.4, 0.7, which approximately corresponds to −26, −20, −14, −8 and −3 dB. In EEG case the lowest SNR tried was 0.1 (−20 dB) because the 0.05 setting usually produced no results. To ensure identical SNRs for all sources, in every simulation run each source RMS amplitude ⟨*s*^2^⟩^1*/*2^ was tuned to obtain sensor the level SNR equal to one of the above numbers.

The following three values of inter source correlations were tried: 0 (uncorrelated sources), 0.5 (moderately correlated sources) and 0.9 (strongly correlated sources). To simplify the analyses all pairwise correlation coefficients between the source time courses were set equal to each other.

As already pointed out the quality of the noise model strongly affects the results for all the discussed methods. We tested three different estimates of the noise covariance with each of them. First matrix ***N*** was calculated using the whole noise dataset with no planted sources; this is referred to as the “*true noise*” setting further on. An estimate of a comparable quality can be obtained in stimuli-driven experiments using “silent” control time intervals with no stimuli present. Second, matrix ***N*** was estimated using 10% of the data at the beginning of the noise dataset (the “*pre-run noise*” setting). This imitated a frequent scenario when noise collection for each subject is performed prior to the experiment itself. In both the MEG and EEG cases such interval was long enough to obtain a robust covariance estimate, as determined by the bandwidth of the data and the number of channels *M* [24]. Note that this approach is much less accurate than the first one due to the noise variability through the entire recording. As a last option, a white diagonal noise was tried (the “*white noise*” setting). This is a frequently used *ad hoc* solution when no noise measurements are available.

A total of 100 simulation runs were conducted per each unique combination of the following independent variables: modality, the number of sources, SNR, inter-source correlations and the noise model. In each such run four localizers: the TRAP MUSIC and SMCMV MAI, MPZ, MER localizers (either SIA MCMV or RAP version) – were tested.

### 3.2. Statistical analyses

Repeated measures ANOVA was applied to look for statistical differences in source localization accuracy and the number of lost sources (the dependent variables). We used a conservative approach where runs with any of the methods failing to find at least one source were excluded from the analyses. Thus occasionally the actual number of runs submitted to ANOVA could be less than 100. If this number was less than 10, this specific combination of independent variables was excluded from consideration. When estimating the average location error for each method in a given run, a maximal common subset of sources found by all four methods was used.

For parameter combinations where statistical differences where detected by ANOVA at 0.05 significance level, post hoc pairwise T-tests between methods were conducted to identify the responsible pairs. Pairwise differences were considered significant at 0.05 level after Bonferroni multiple comparisons correction (a factor of 6).

### 3.3. Comparisons with MIA

A separate set of simulations was conducted to compare SMCMV localizers with MIA MCMV algorithm [18]. Due to relatively long run times for MIA, we limited the analyses to the EEG case and chose a single best performing beamformer type, that is MER, to look at. Knowing how the latter compares to the TRAP MUSIC [7], one can also make conclusions about MIA versus TRAP performance. Again 100 simulation runs were done per each unique combination of the number of sources, SNR value, inter-source correlation and the noise model, and the same statistical analyses were applied.

## 4. Results

Due to space limitations we present only a subset of results, for a specific case when RAP beamformer solutions (9) were substituted for *P* in equation (10). More results, in particular for SIA MCMV option, can be found in *Supplemental materials*. The presentation is structured in accordance with the noise model used, as the latter plays a key role in how all the methods perform.

### 4.1. True noise

As should be expected, both the TRAP MUSIC and SMCMV performed best in this case. On average for both modalities all techniques achieved maximum possible localization accuracy (mean error less than 3 mm given the measurement grid step size 5 mm) in most cases (see Fig. 1 for MEG and Fig. S.1 in *Supplemental materials* for EEG data). Larger location errors occurred for the lowest SNR values (less than 0.2, or -14 dB) for *n* = 10, 20 and especially for strong source correlations. For the latter case SMCMV power localizers MAI and MPZ practically matched the TRAP MUSIC results. MER accuracy was marginally worse than TRAP for MEG data reaching statistical significance at *n* = 10, 20 and *C* = 0.9; no such effect was observed for EEG (Fig. S.1).

**Figure 1:**
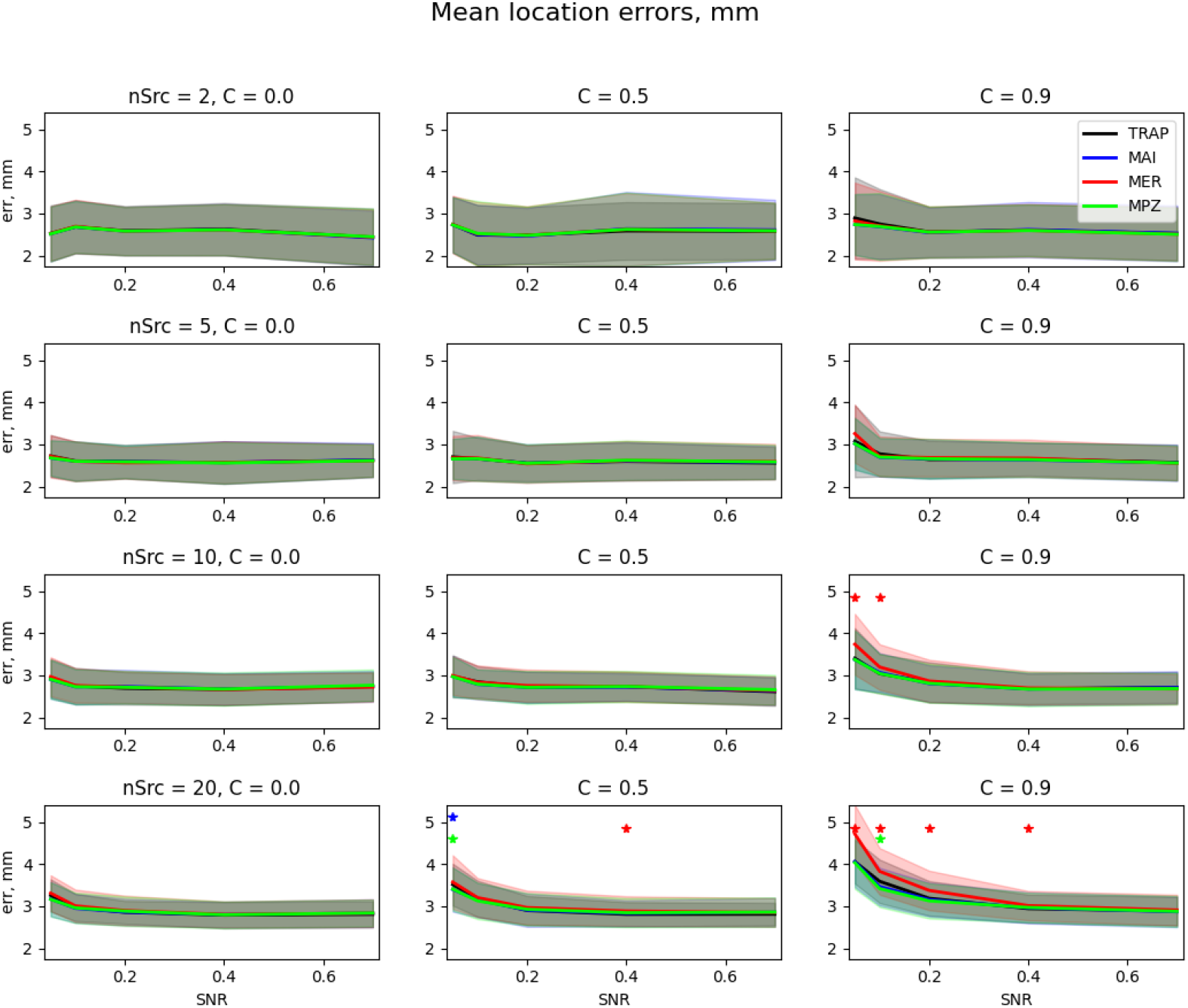
Average source localization errors for **TRAP MUSIC**(black) and **SMCMV RAP beamformer** localizers MAI (blue), MPZ (green) and MER (red) as functions of SNR, depending on the number of sources and inter-source correlations in the **MEG true noise** case. The shadowed areas mark one standard deviation around corresponding curves. Asterisks denote SNR values where statistical differences between the respective SMCMV method and TRAP MUSIC were significant. In this particular case all SMCMV results practically coincided so MAI and MER lines are mostly obscured by the MPZ line.

The same situation was observed with the number of missed sources (see Fig. S.2, S.3). Typically all sources were found in most runs, performance of all the methods being approximately the same. Probability of missing a source naturally increased with their total number *n*. An average number of lost sources exceeded 1 only for MER applied to MEG data, in an extreme case *n* = 20*, C* = 0.9 (Fig. S.2).

Overall, for all practical purposes in the true noise case SMCMV provided no advantages over the TRAP MUSIC approach.

### 4.2. Pre-run noise

In this case we were dealing with a much cruder noise model than in the previous section, and the results were quite different (Fig. 2, 3). For lower SNRs and especially for higher inter-source correlations, SMCMV beamformers demonstrated significantly better spatial accuracy than the TRAP MUSIC. For very high SNRs accuracy of all methods became approximately equal tending to the limit dictated by the measurement grid step. Qualitatively the picture looked the same for both MEG and EEG cases, but for MEG the differences with MUSIC reached statistical significance more often due to its higher spatial resolution. The evoked SMCMV beamformer (MER) provided the best results for both modalities and in MEG case attained maximum possible accuracy for all parameter combinations. As for the lost sources (see Fig. 4, and Fig. S.4 in *Supplemental materials*), SMCMV localizers missed less sources than TRAP in all cases, and the differences were statistically significant for wide range of SNRs and source correlations. Again MER demonstrated the most improvements especially for low SNR values, where power localizers MAI and MER did not show big changes compared to TRAP MUSIC (Fig. 4).

**Figure 2:**
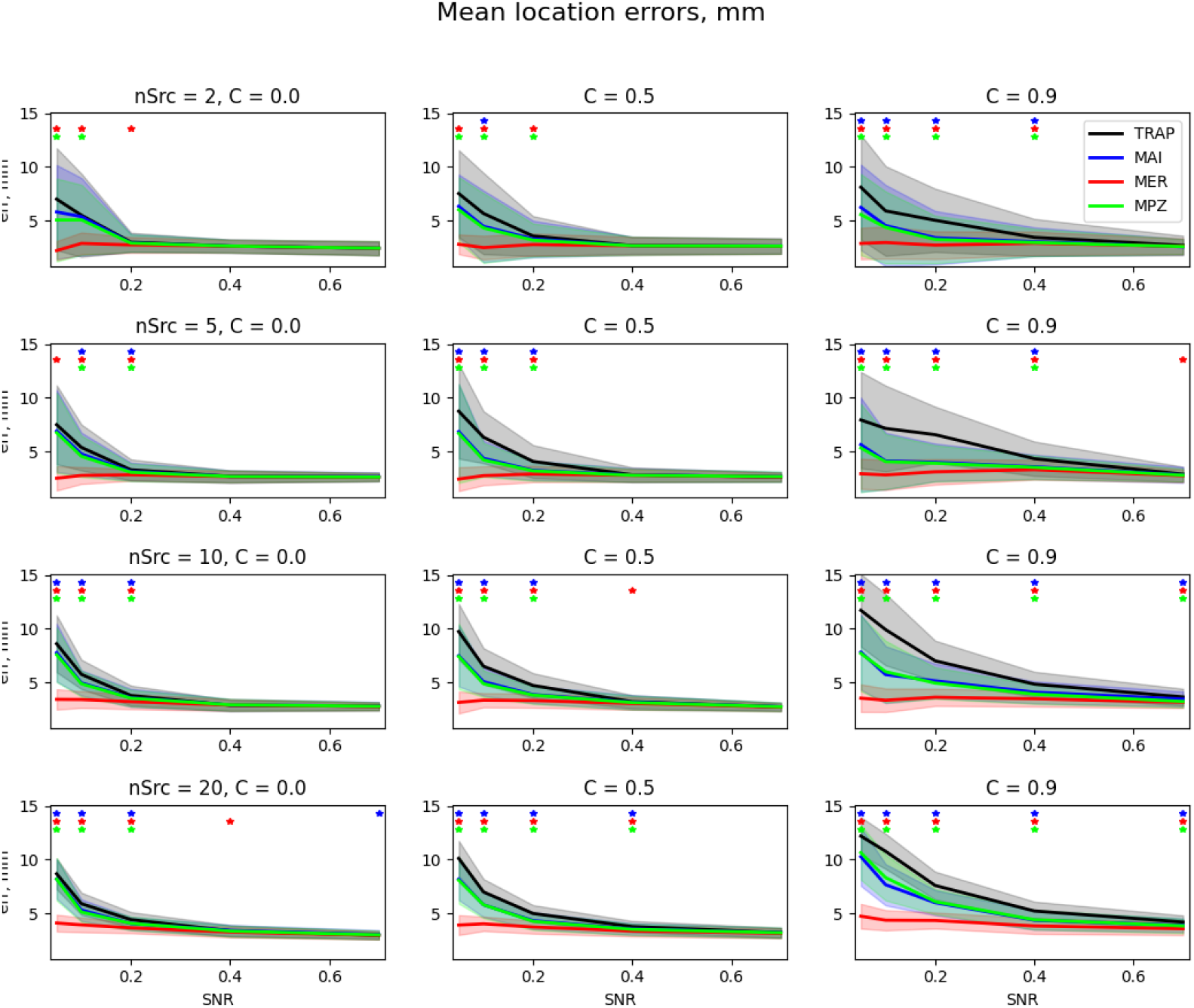
Average source localization errors for **TRAP MUSIC** and **SMCMV RAP beamformer** in **MEG pre-run noise** case. The notation is the same as in Fig. 1.

**Figure 3:**
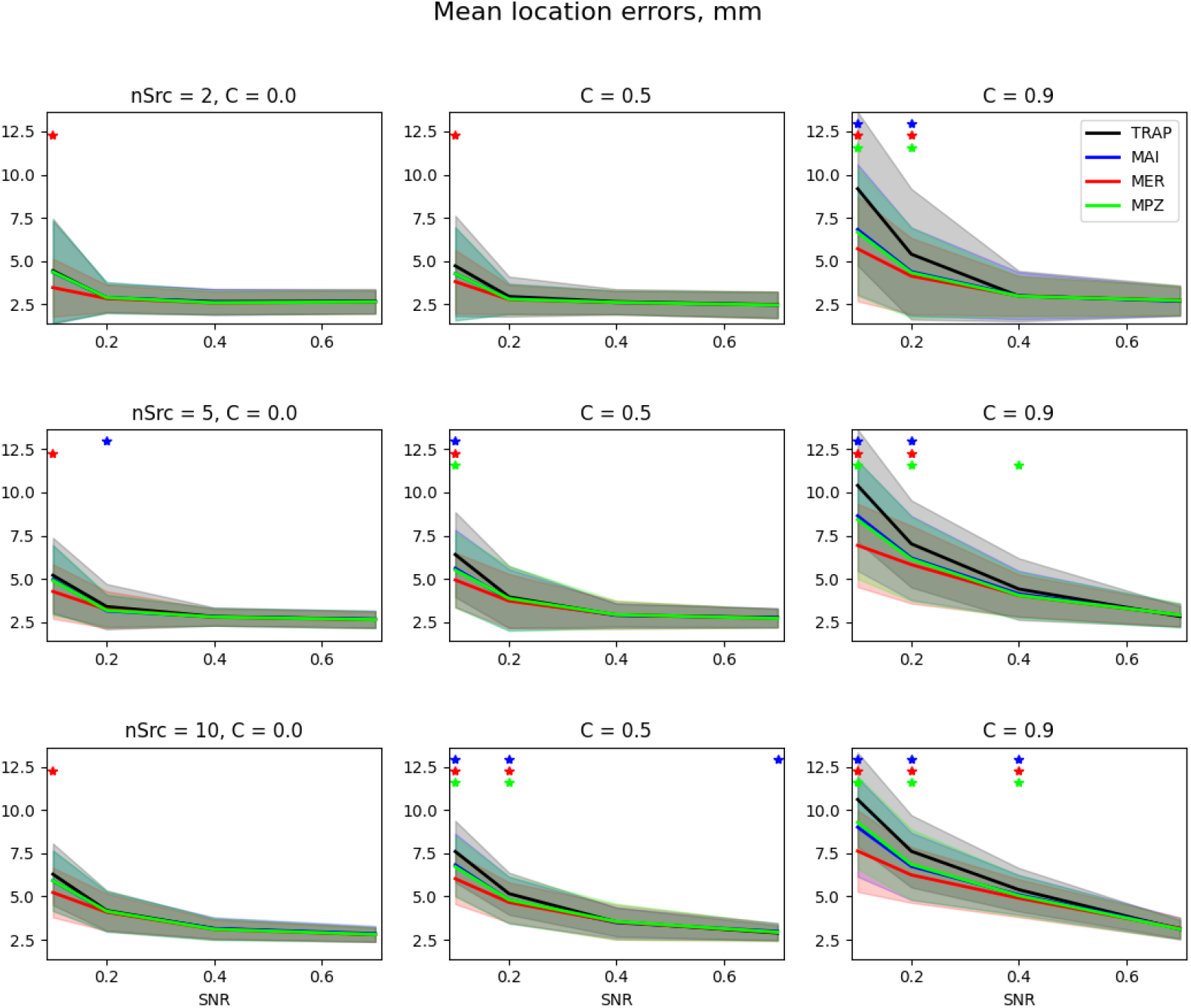
Average source localization errors for **TRAP MUSIC** and **SMCMV RAP beamformer** in **EEG pre-run noise** case. The notation is the same as in Fig. 1.

**Figure 4:**
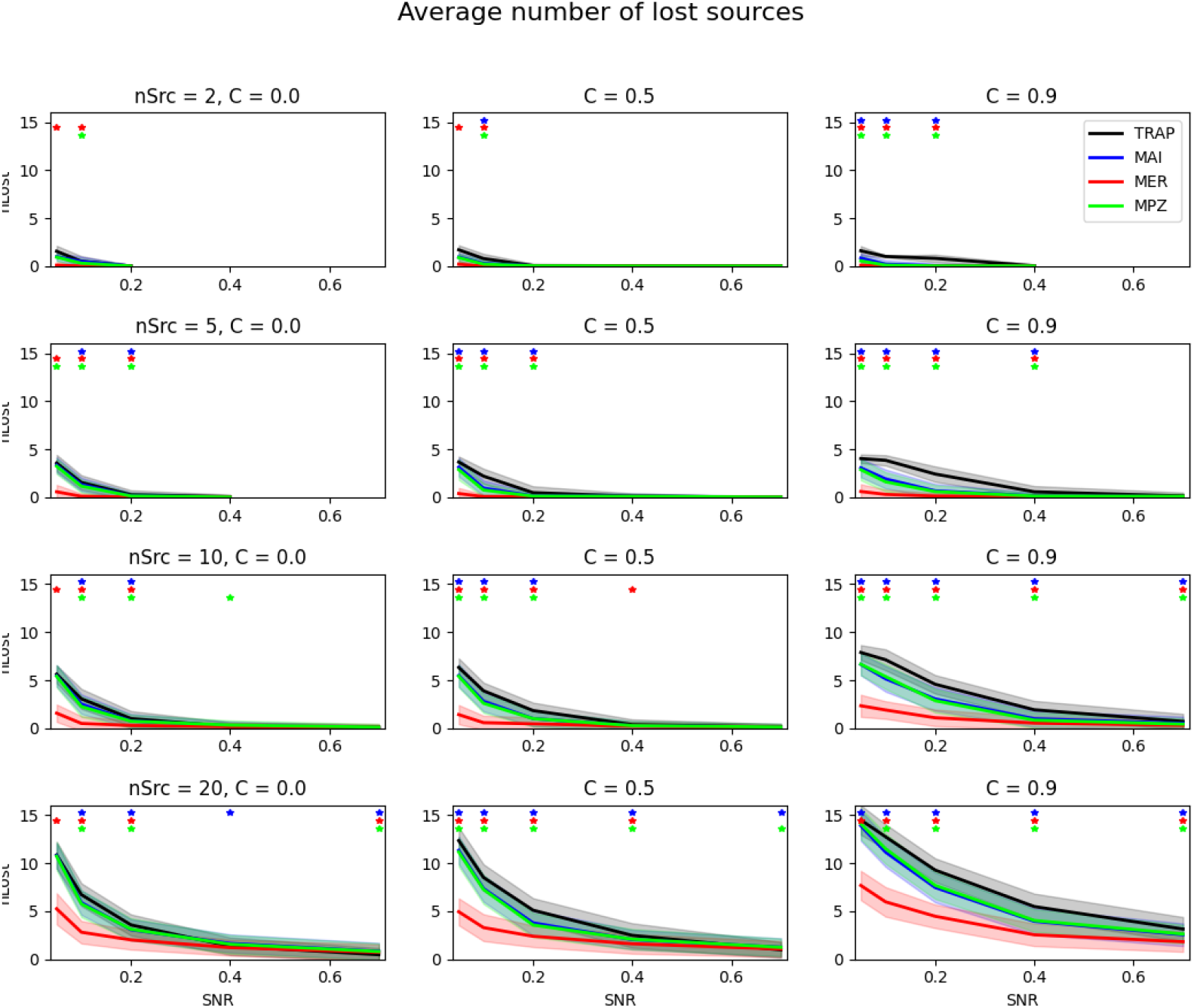
Average numbers of lost sources for **TRAP MUSIC** and **SMCMV RAP beamformer** localizers in the **MEG pre-run noise** case. The notation is the same as in Fig. 1.

In summary, in the pre-run noise case SMCMV proved to be better than TRAP MUSIC from very low to moderate SNR values. Improvements were especially significant for larger number of sources and stronger inter-source correlations.

### 4.3. White noise

As could be expected, performance of all methods in this case was much worse than for the true or pre-run noise (see Fig. 5, 6). Manifold increase in average localization errors compared to the pre-run noise case was observed, and for higher correlations (*C* = 0.9) even large source SNRs did not necessary result in significant increase in accuracy, especially for the TRAP MUSIC solution. The MER (evoked) solution was the best again but now clear differences in performance among the power-based methods was observed also, especially for EEG and for higher correlations, with the TRAP MUSIC being the less accurate and MPZ being the most accurate of all.

**Figure 5:**
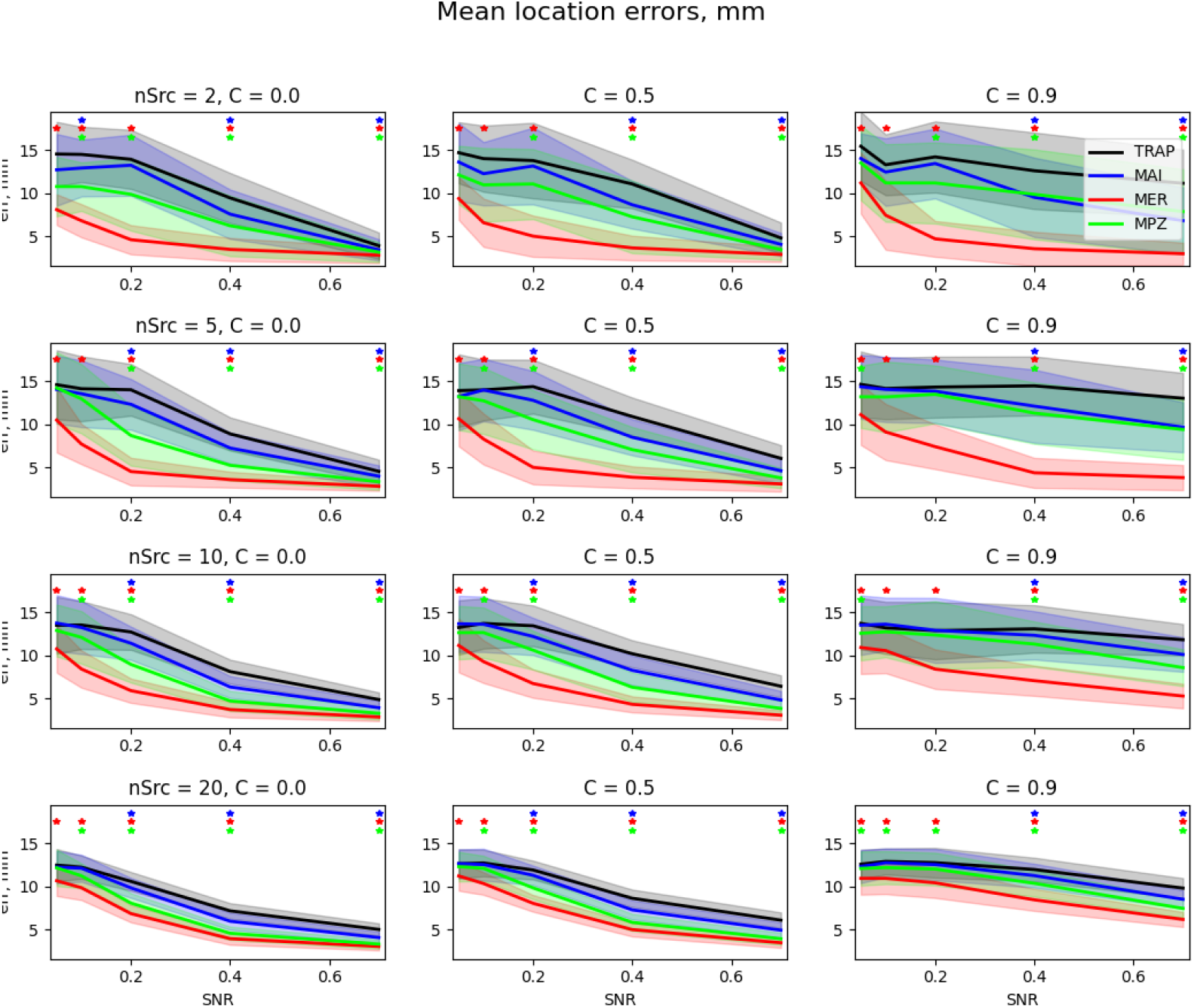
Average source localization errors for **TRAP MUSIC** and **SMCMV RAP beamformer** localizers in the **MEG white noise** case. The notation is the same as in Fig.1.

**Figure 6:**
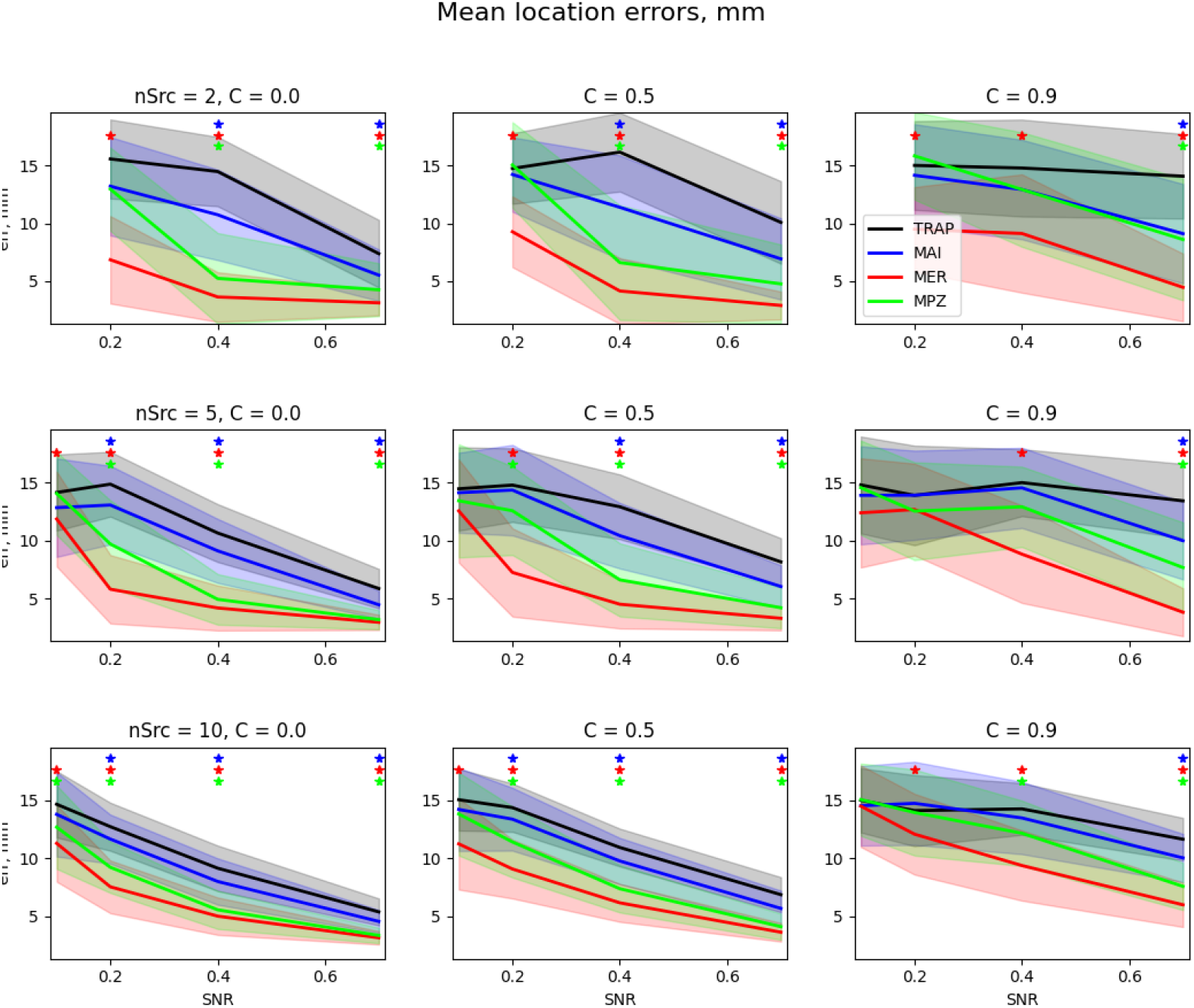
Average source localization errors for **TRAP MUSIC** and **SMCMV RAP beamformer** localizers in the **EEG white noise** case. The notation is the same as in Fig.1.

As for the missed sources, the situation was the worst for the TRAP MUSIC again because it solely relied on correct source and noise subspace identification obviously missing in this case. For *n* = 2, 5 and the lowest considered SNR values TRAP MUSIC did not find any sources in most runs for both EEG and MEG cases; for *n* = 10 typically 80% of sources were lost (see Fig. S.5, S.6). With growing SNRs more sources were found especially for *C* = 0 and *C* = 0.5 but not necessarily for *C* = 0.9. SMCMV power localizers did noticeably better especially in EEG case, while the evoked SMCMV MER localizer provided the most significant improvements.

### 4.4. Comparisons with MIA MER beamformer

Fig. 7 shows results of running MIA MER and RAP SMCMV MER on the same 100 simulated EEG datasets using the pre-run noise estimate. One can see that MIA exhibited smaller localization errors for lower SNRs and *n* = 2, 5 when correlations were high (*C* = 0.9). SMCMV MER showed approximately the same accuracy as MIA for *n* = 2 and better accuracy for larger number of sources (*n* = 5, 10) and *C* = 0, 0.5. For *n* = 10, *C* = 0.9 SMCMV outperformed MIA for higher SNR values.

**Figure 7:**
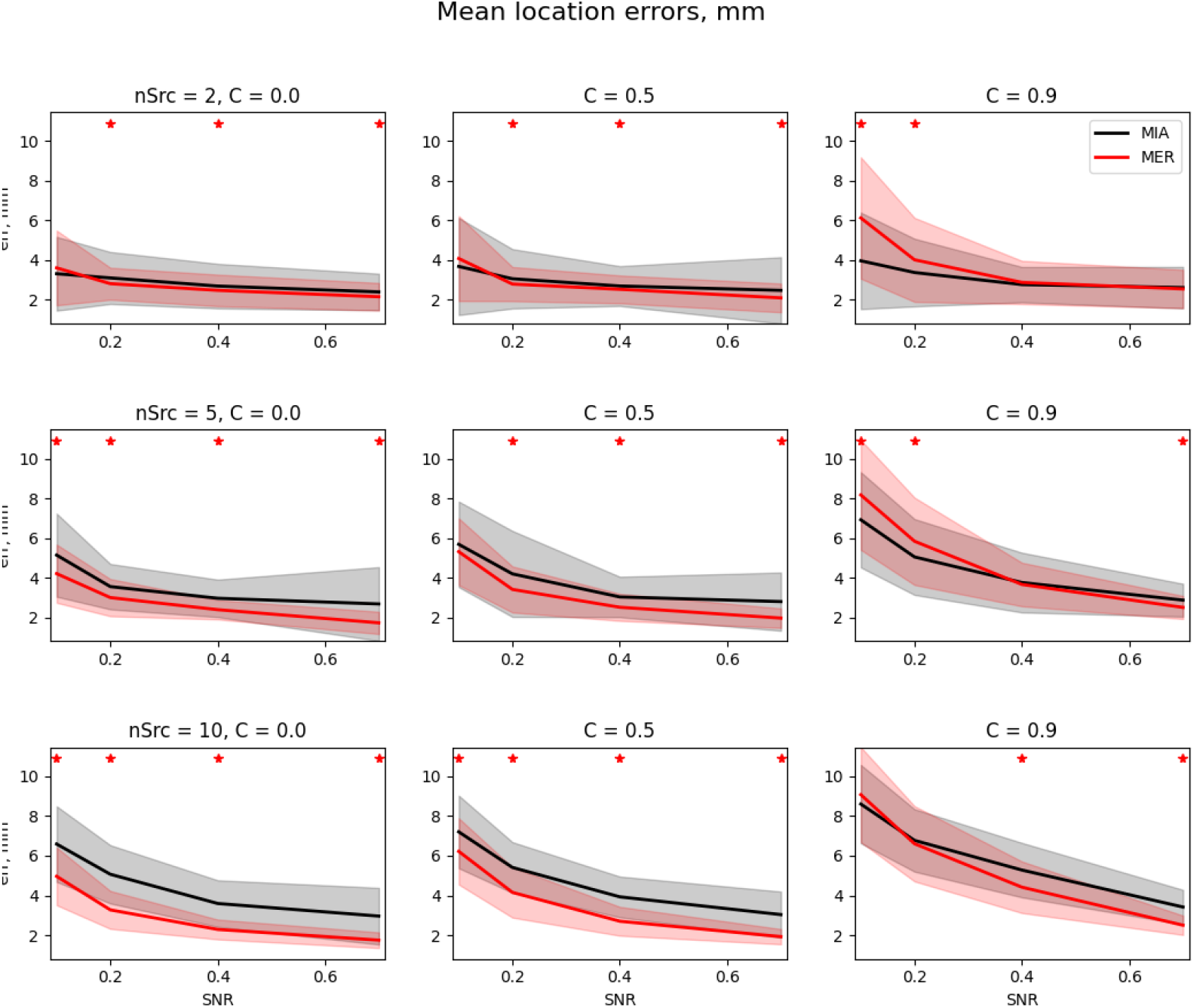
Average source localization errors for **MCMV MIA**(black) and **SMCMV RAP beamformer MER**(red) localizers in the **EEG pre-run noise** case. Other notation is analogous to Fig.1.

Note however that reported localization errors only refer to the subset of sources found by *both* methods in any given run. When one looks at the total numbers of sources missed by either method (Fig. 8), bigger differences become evident. It turns out that the results were close only for small *n*’s and small SNR values. In other cases MIA missed many more sources than SMCMV MER not only for larger *n* but also with *growing* SNRs and *decreasing* source correlations. This is in contrast to SMCMV MER whose behavior was exactly the opposite: for high enough SNRs it found all sources in the majority of the runs. The situation in other cases (power beamformers, MEG, true noise, etc) was qualitatively the same. Thus one can conclude that overall forward SMCMV did a better job than MIA MCMV for a wide range of parameter values, especially with respect to the total number of found sources.

**Figure 8:**
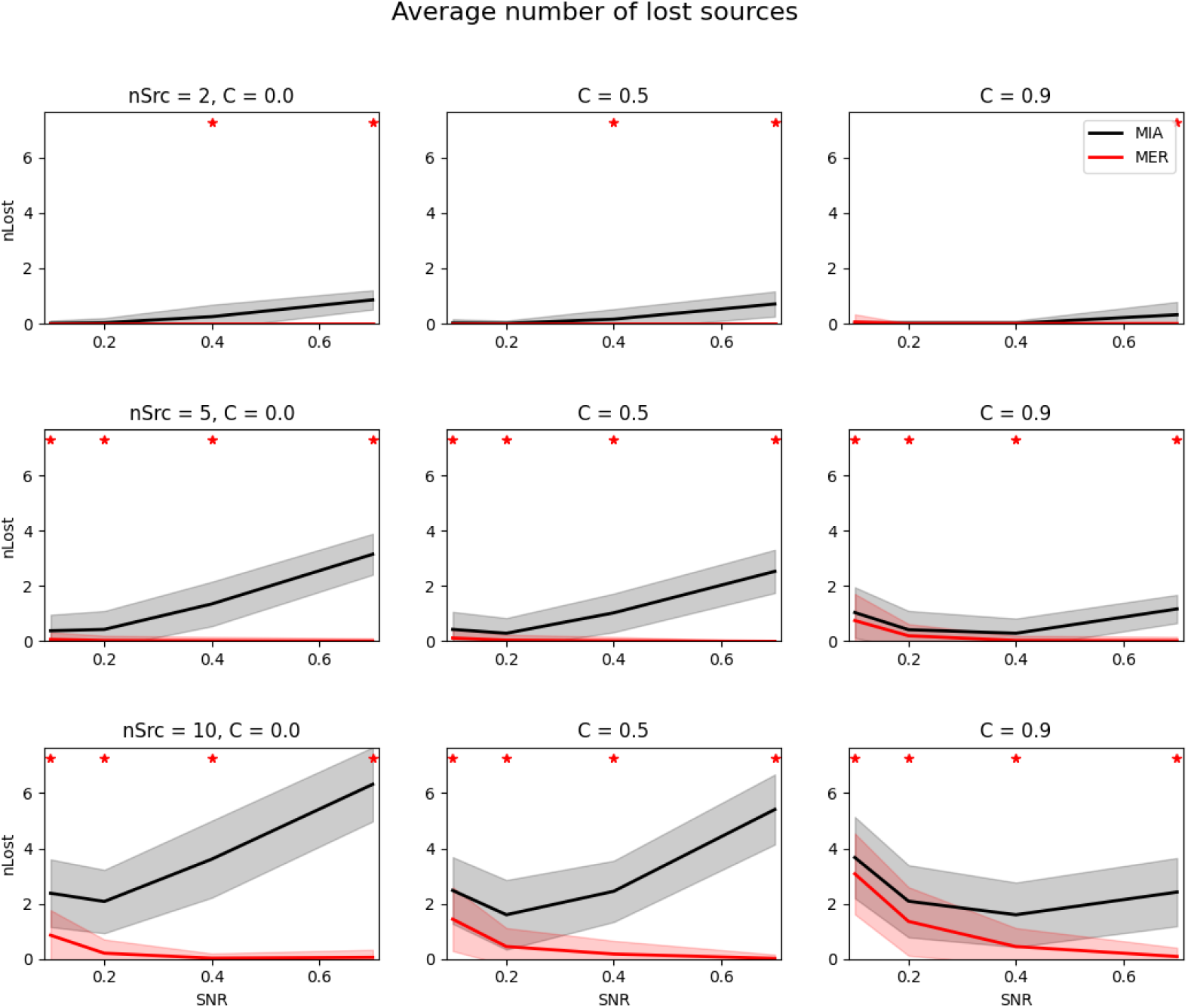
Average numbers of lost sources for **MCMV MIA**(black) and **SMCMV RAP beamformer MER**(red) localizers in the **EEG pre-run noise** case. Other notation is analogous to Fig.1.

## 5. Discussion

In this work, we introduced a family of multi-source localizer functions with the following attractive properties: they are *unbiased* in error-free ideal situations irrespective to the SNR values or source correlations; they come with a simple and computationally inexpensive way of finding all the sources; and they demonstrate robust performance and good accuracy when realistic errors are present. This is achieved by combining the RAP MUSIC and multisource beamformer methods to construct localizer functions incorporating advantages of either technique.

The simple SMCMV algorithm suggested above is not the only one possible. In particular, in contrast to the TRAP MUSIC [7] the generic SMCMV localizers defined by Eq.(10) are calculated without truncating the source subspace at each iteration step (see section *RAP and TRAP MUSIC* for details). Such truncation can be easily included but is unnecessary as simulations confirm, because residuals of the out-projected sources have negligible SNRs and by design do not produce false maxima of SMCMV localizers. Also, more sophisticated “combined” approaches could be constructed using multiloop iterations both on the MUSIC side as in [25] and on the MCMV side as with MIA [18]. However we believe that while possibly yielding improvements in accuracy in certain situations, a manifold increase in computational times would generally outweigh the benefits. Besides simple and computationally effective inverse solutions are more preferable when it comes to large group studies and automated analyses. This is where SMCMV localizers fit well because they are no more sophisticated than RAP MUSIC or traditional single source beamformers.

Results presented in the previous section refer to a particular case when the RAP beamformer expressions (9) were substituted for *P* in Eq. (10). When SIA SMCMV (7) were used instead the results could be marginally better or worse depending on the situation. Specifically we found that for the evoked (MER) SMCMV the RAP beamformer version typically performed the same or better than the SIA version, while for power localizers either one could be a touch better or worse but generally did not differ significantly for all practical purposes (see Fig. S.7, S.8 in *Supplemental materials*). Note also that while it is natural to substitute a known localizer for *P* (***H***; ***r***) in (10), this is not a requirement (for that matter *P* does not need to be a localizer at all). For example it turned out that replacing pseudoinverses 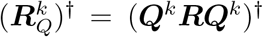 everywhere in the RAP beamformer part of SMCMV with ***Q**^k^**R***^−1^***Q**^k^* marginally improved localization accuracy for MER (Fig. S.9). In any case, more research is needed to identify optimal choices for *P* depending on practical situations.

In this study we compared SMCMV with TRAP MUSIC and MIA as representatives of MUSIC and beamformer families of the inverse solutions. As both are extensively studied, one can also judge how SMCMV compares to other techniques. For example, SMCMV should do better than single source beamformers and single loop multi-source beamformers (like SIA or RAP beamformer) in most situations. At the same time just like any MUSIC or beamformer solution, SMCMV is only applicable for identifying low rank activations and is not suitable for analysing distributed brain activity. Alternative techniques such as minimum norm estimation are more appropriate in the latter case.

Our computer simulations once again demonstrated importance of an accurate noise covariance estimate for both MUSIC and beamformer solutions. With the true noise model and proper pre-whitening of the data TRAP MUSIC did an excellent job even for small SNRs (0.2, or -14 dB) and high correlations (0.9), and it was hard to do better (Fig. 1). However the “true noise” estimate is rarely possible in real measurements. When the quality of the noise approximation deteriorated so did the accuracy of the TRAP MUSIC localizer, especially with increasing source correlations. While SMCMV results demonstrated the same trend they now outperformed the TRAP MUSIC significantly (Fig. 2 - 6).

We also want to notice performance of the SMCMV evoked (MER) localizer. In a single occasion when the noise model was precise and the TRAP MUSIC was at its best, MER showed slightly *worse* results in the MEG case for low SNRs and high correlations (Fig. 1). In all other circumstances SMCMV MER provided big improvements compared not only to the TRAP MUSIC but to the SMCMV power localizers as well (see Fig. 2 - 6 and other results in *Supplemental materials*). Overall, we found MER SMCMV to be by far the best option among all tested for the evoked scenarios, both with respect to the localization accuracy and to the total numbers of successfully identified sources.

Using the evoked case as an example, our comparisons of SMCMV with MIA MCMV algorithm proved that either one could have better localization accuracy depending on the parameter combination, if measured using a subset of sources simultaneously found by both techniques (see Fig. 7 and discussion in section *Comparisons with MIA MER beamformer*). However in many situations, especially with increasing SNRs MIA was missing many sources, up to 50% for highest SNR values. Nothing like that happened with SMCMV which typically identified more sources and tended to find them all at higher SNRs. A most likely explanation is that for high SNRs MIA iterations were stuck in local extrema of the localizer function in the parameter space. This did not happen for SMCMV due to the out-projection mechanism built into the MUSIC component of the cost function.

## 6. Conclusion

In this study we presented a novel set of localizer functions for identifying sources of EEG and MEG brain activity in case of low rank focal activations. They combine the MUSIC approach based on the signal and noise subspace separation, and a multiple constrained minimum variance (MCMV) spatial filter approach. The localizers come with a computationally efficient algorithm, which is analytically shown to yield precise locations and orientations of all sources for arbitrary values of signal to noise ratios and inter-source correlations when no measurement or modeling errors are involved.

We validated the SMCMV method by extensive computer simulations involving measured human EEG and MEG data and realistic lead fields modeling errors, for different numbers of sources and a range of values for SNR and inter-source correlations, and compared it to the TRAP MUSIC and MIA MCMV results.

It was found that when the true noise model was not available, which is the case for most practical situations, SMCMV localizers typically outperformed TRAP MUSIC across a wide range of parameter combinations.

Comparisons with MIA MCMV showed that MIA beamformer provided more accurate localizations when the number of sources was small (*n* = 2, 5), SNR was low (less than −8 dB) and the correlations were strong (*C* = 0.9). SMCMV was better in most other cases.

Overall, we conclude that SMCMV provides a relatively simple and reliable way of localizing brain sources in MEG and EEG, and achieves significant improvements over both RAP/TRAP MUSIC and MCMV beamformer in a variety of situations, especially for larger numbers of sources and high inter-source correlations. Due to its computational efficiency SMCMV is well suited for group studies involving processing of large amounts of neuroimaging data.

## Appendix A. Unbiased property of SMCMV localizers

Start with the MUSIC localizer *μ*(***h***), where ***h***(***r**, **o***) is a lead field of a dipole source at test location ***r*** with orientation ***o***: ***h*** = ***L***(***r***)***o*** (see section 2.2). *μ*(***h***) represents a squared ratio of the length of the projection of ***h*** on the source subspace ***U***, equal to ***UU** ^T^ **h***, to the length of ***h***:

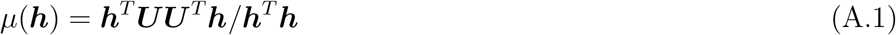

When source orientation ***o*** is not known, we take the one that maximizes *μ* given ***r***, which leads to Eq.(4).

Differentiating (A.1) with respect to ***h*** we get *dμ* = 2(***h**^T^ **h***)^−2^***h**^T^* (***hh**^T^ **UU** ^T^* − ***UU** ^T^ **hh**^T^*)*d**h***. When ***h*** ∈ ***U*** we have ***UU** ^T^ **h*** ≡ ***h*** therefore expression in brackets becomes zero. Thus *dμ*(***h***) = 0 when ***h*** belongs to the signal subspace (the unbiased property of the MUSIC localizer). The same proof holds when ***UU** ^T^*, ***h*** in (A.1) are replaced with their “RAP” versions (*Q^k^U*)(*Q^k^U*)^†^, *Q^k^h* because from *h* ∈ *U* follows *Q^k^h* ∈ *Q^k^U*, therefore at every RAP iteration step *dμ*(***h**^k^*^−1^; ***r***) = 0 when corresponding test lead field ***h**^k^*(***r***) ∈ ***U***.

The inverse of a SMCMV localizer (10) is

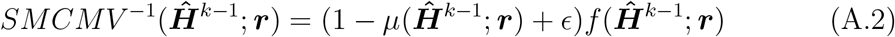

with 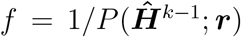 being a smooth limited positive function. Differentiating (A.2) with respect to ***h*** we have *d*(*SMCMV* ^−1^) = −*fdμ* + (1 − *μ* + *ϵ*)*df* → 0 when ***h**^k^* ∈ ***U**, ϵ* → 0 because in this case *dμ*(***h**^k^*) = 0 and *μ*(***h**^k^*) = 1. Thus *SMCMV* ^−1^ has extremum when ***h**^k^* ∈ ***U***, which is a minimum equal to 0 as follows from (A.2). Equivalently expression (10) reaches a local maximum, which proves the unbiased property of SMCMV localizers.

## Appendix B. Surrogate brain noise data

The M/EEG data collected from a real subject even in a resting state condition may still contain occasional synchronous events or reflect intrinsic connectivity. In computer simulations one wants to create an environment where the activity of interest is represented entirely by the modeled brain sources without any uncontrolled contributions from the brain noise background. To achieve this the following randomization procedure is applied to the data.

First, the original data ***b***(*t*) in the sensor space is expanded in principal components (PC) basis:

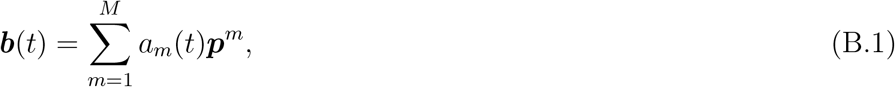

where PC vectors ***p**^m^* are the eigenvectors of the covariance matrix ***R***. Functions *a_m_*(*t*) = ***b**^T^* (*t*)***p**^m^* represent PC time courses. Second, for each *a_m_*(*t*) we create its randomized version *α_m_*(*t*) by calculating their Fourier transforms *r_mω_* exp(*iϕ_mω_*) = exp(*iωt*)*a_m_*(*t*)*dt*, randomizing phases 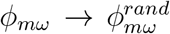 and then performing the inverse Fourier transformation:

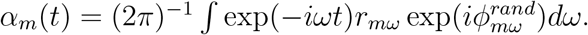

Finally, we obtain the surrogate brain noise data ***b**^s^*(*t*) by replacing the original PC time courses *a_m_*(*t*) in (B.1) with *α_m_*(*t*):

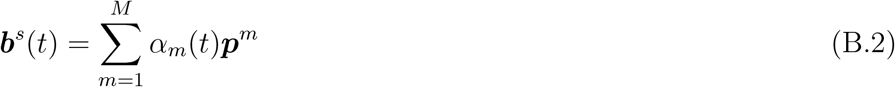

Due to the phase randomization any synchronous activity that existed in the original data is obliterated. At the same time, both the power spectra and the covariance of ***b***(*t*) are preserved.

Indeed, using Eq.(B.1) for the Fourier spectrum *b_kω_* of any sensor channel *k* = 1*, …, M* we get 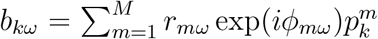, therefore its power spectrum is 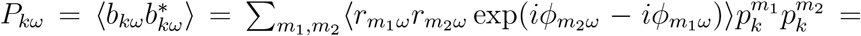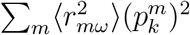. The non-diagonal terms *m*_1_ ≠ *m*_2_ canceled out because PC time courses as well as their spectra are uncorrelated. However the power spectrum of the surrogate data for the same channel 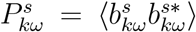 calculated using Eq.(B.2) yields exactly the same result, because now the nondiagonal terms are canceled due to the phase randomization and the diagonal ones are still equal to 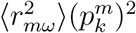.

Similarly, the covariance matrix ***R**^s^* of the surrogate data equals to ***R*** (see (B.2)): 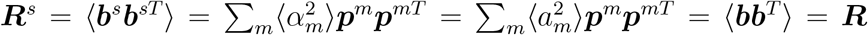 due to 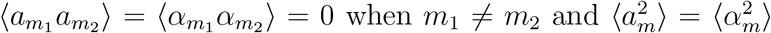 for any *m* = 1*, …, M*.

## Appendix S. Supplemental materials

**Figure S.1:**
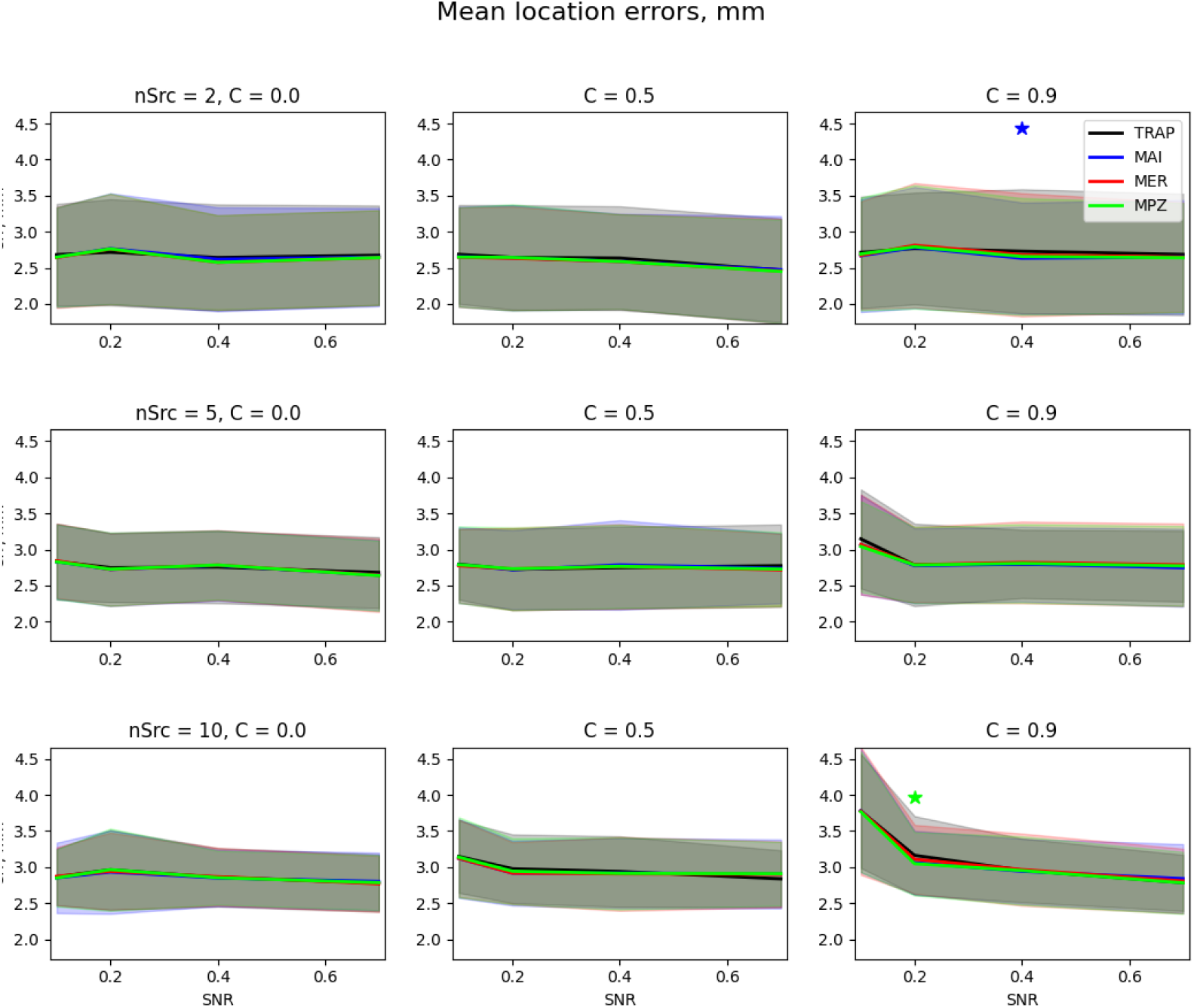
Average source localization errors for **TRAP MUSIC**(black) and **SMCMV RAP beamformer** localizers MAI (blue), MPZ (green) and MER (red) as functions of SNR, depending on the number of sources and inter-source correlations, for the **EEG true noise** case. The shadowed areas mark one standard deviation around corresponding curves. Asterisks denote SNR values where statistical differences between respective SMCMV method and TRAP MUSIC were statistically significant.

**Figure S.2:**
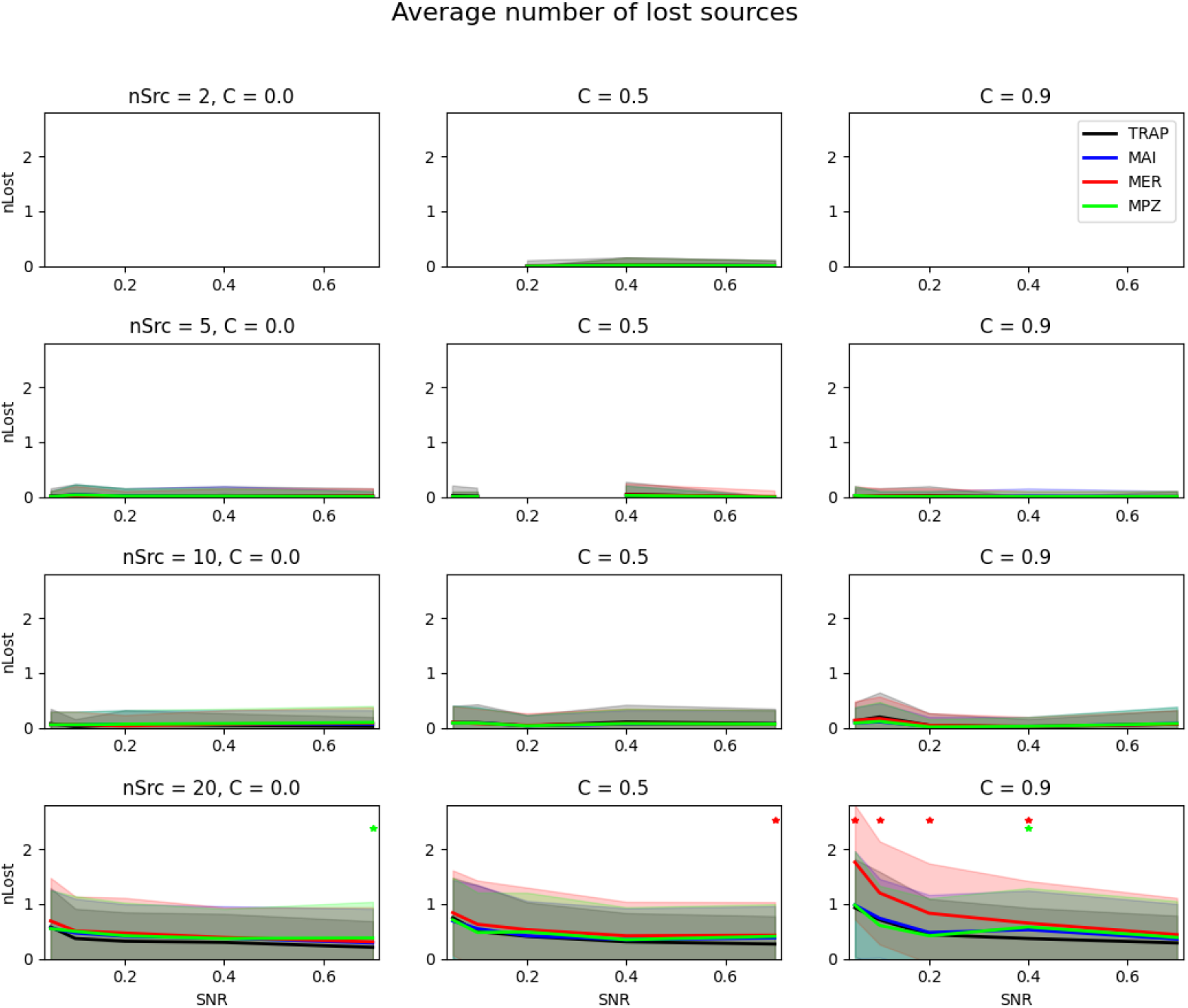
Average number of lost sources for **SMCMV RAP beamformer** localizers in the **MEG true noise** case. The notation is the same as in Fig. S.1

**Figure S.3:**
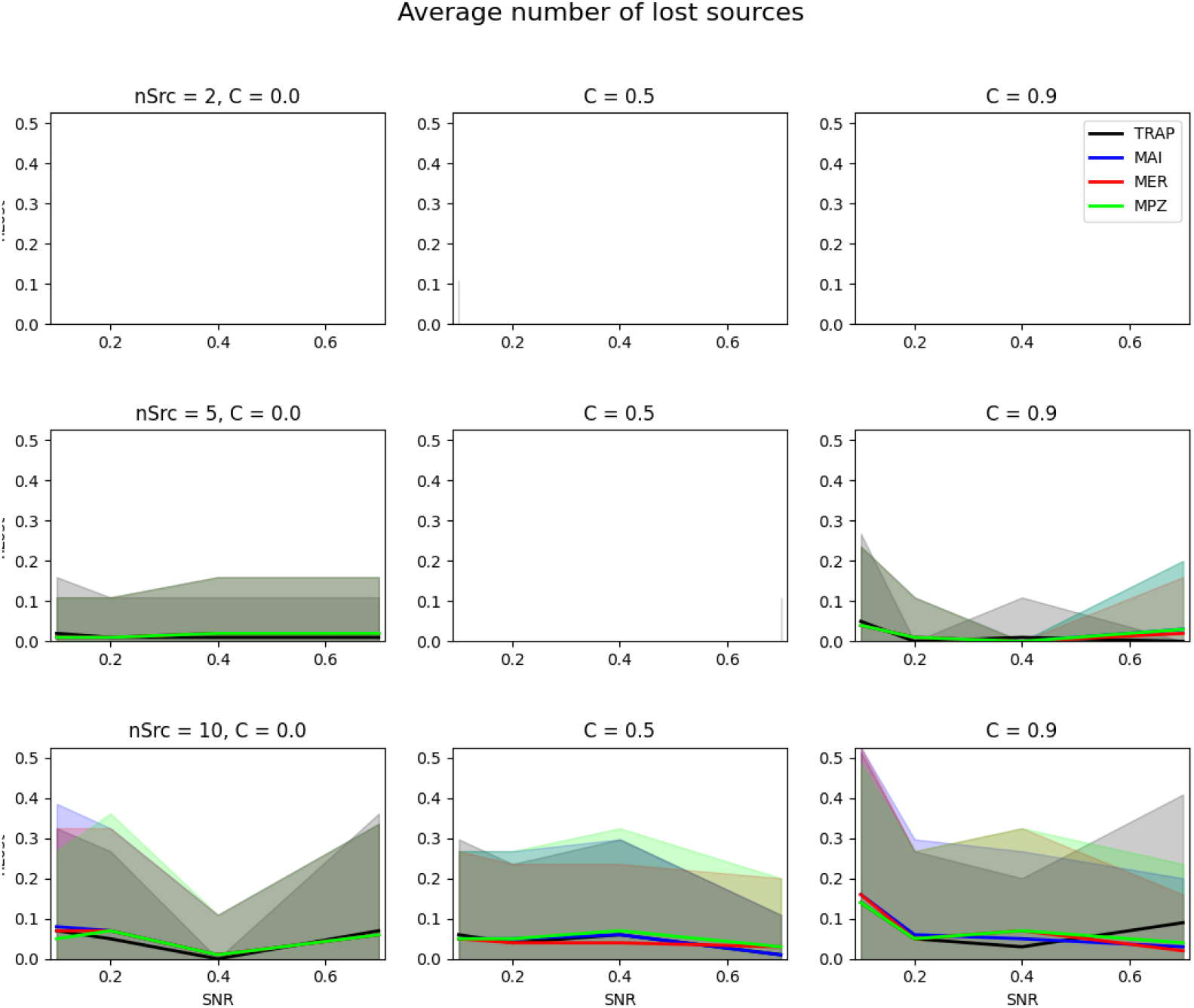
Average number of lost sources for **SMCMV RAP beamformer** localizers in the **EEG true noise** case. The notation is the same as in Fig. S.1

**Figure S.4:**
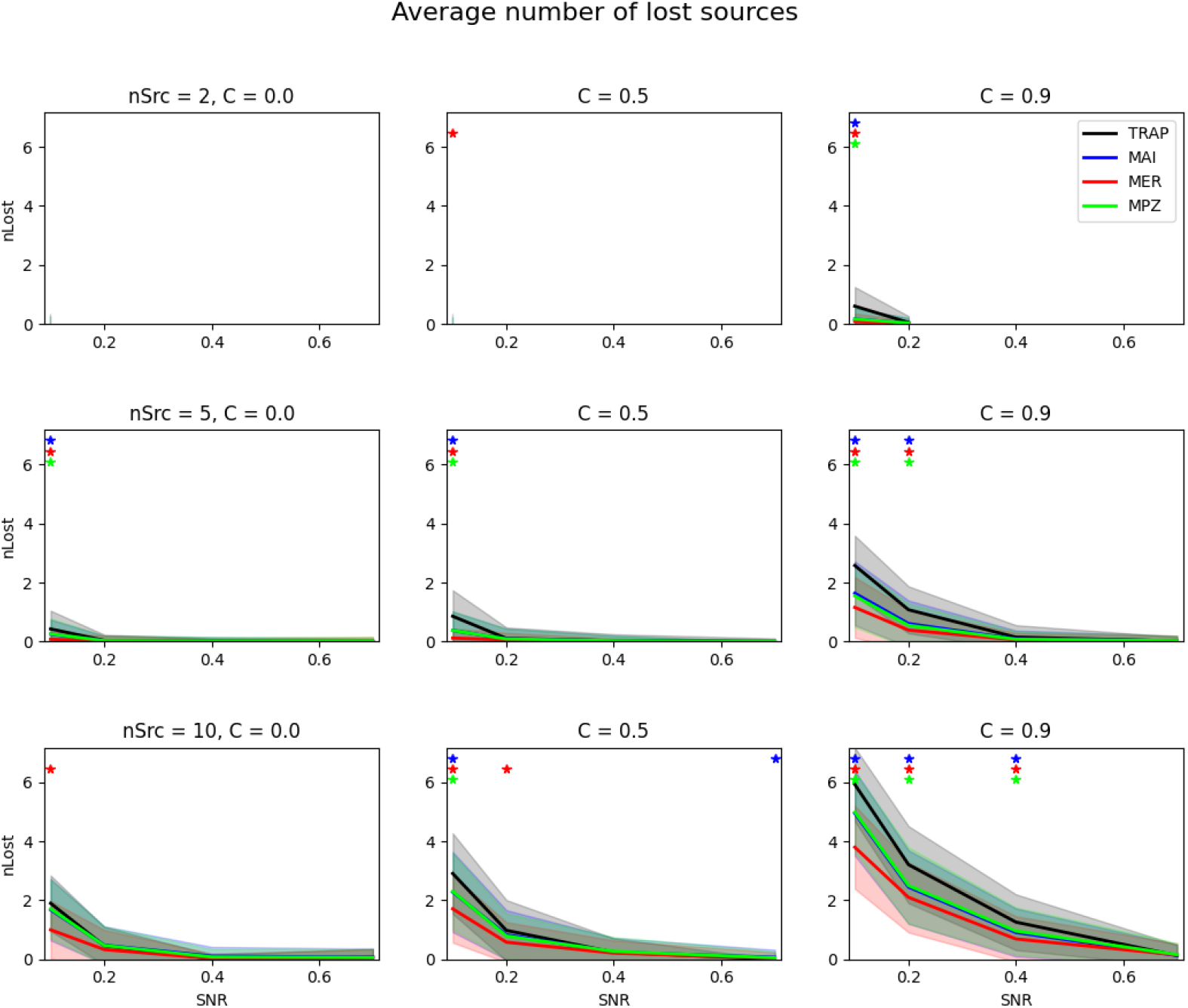
Average numbers of lost sources for **TRAP MUSIC** and **SMCMV RAP beamformer** localizers in the **EEG pre-run noise** case. The notation is the same as in Fig. S.1.

**Figure S.5:**
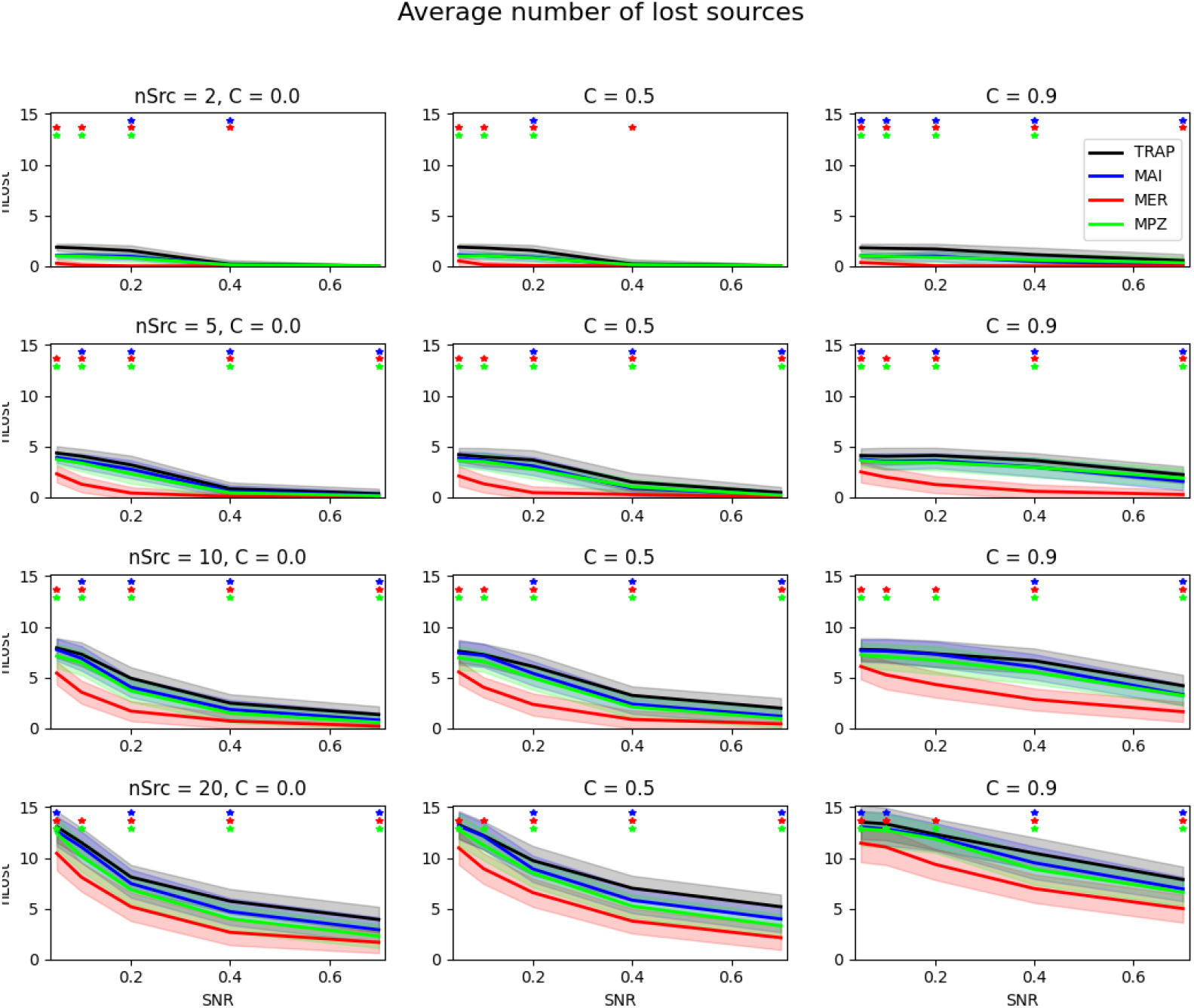
Average numbers of lost sources for **TRAP MUSIC** and **SMCMV RAP beamformer** localizers in the **MEG white noise** case. The notation is the same as in Fig. S.1.

**Figure S.6:**
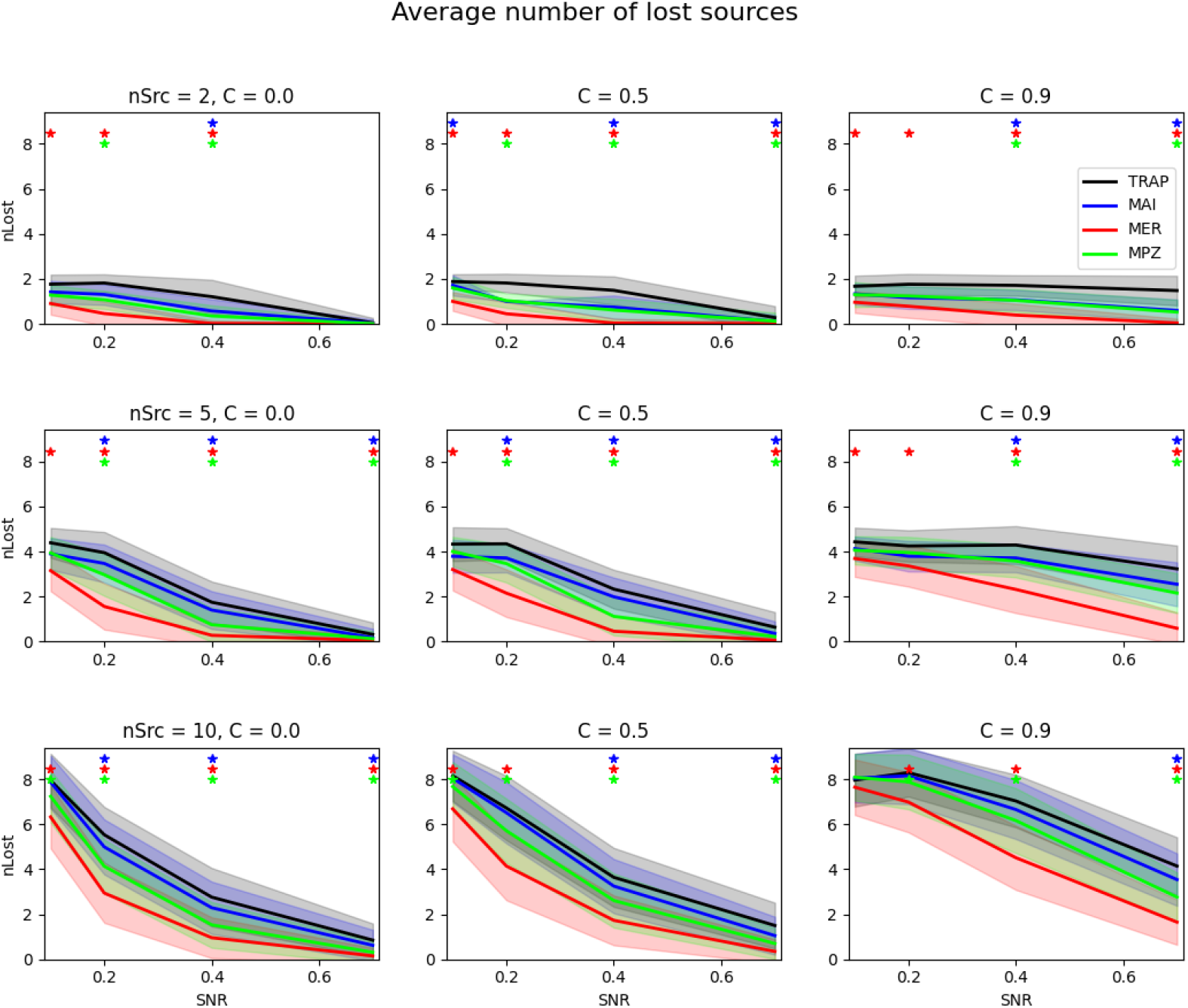
Average numbers of lost sources for **TRAP MUSIC** and **SMCMV RAP beamformer** localizers in the **EEG white noise** case. The notation is the same as in Fig. S.1.

**Figure S.7:**
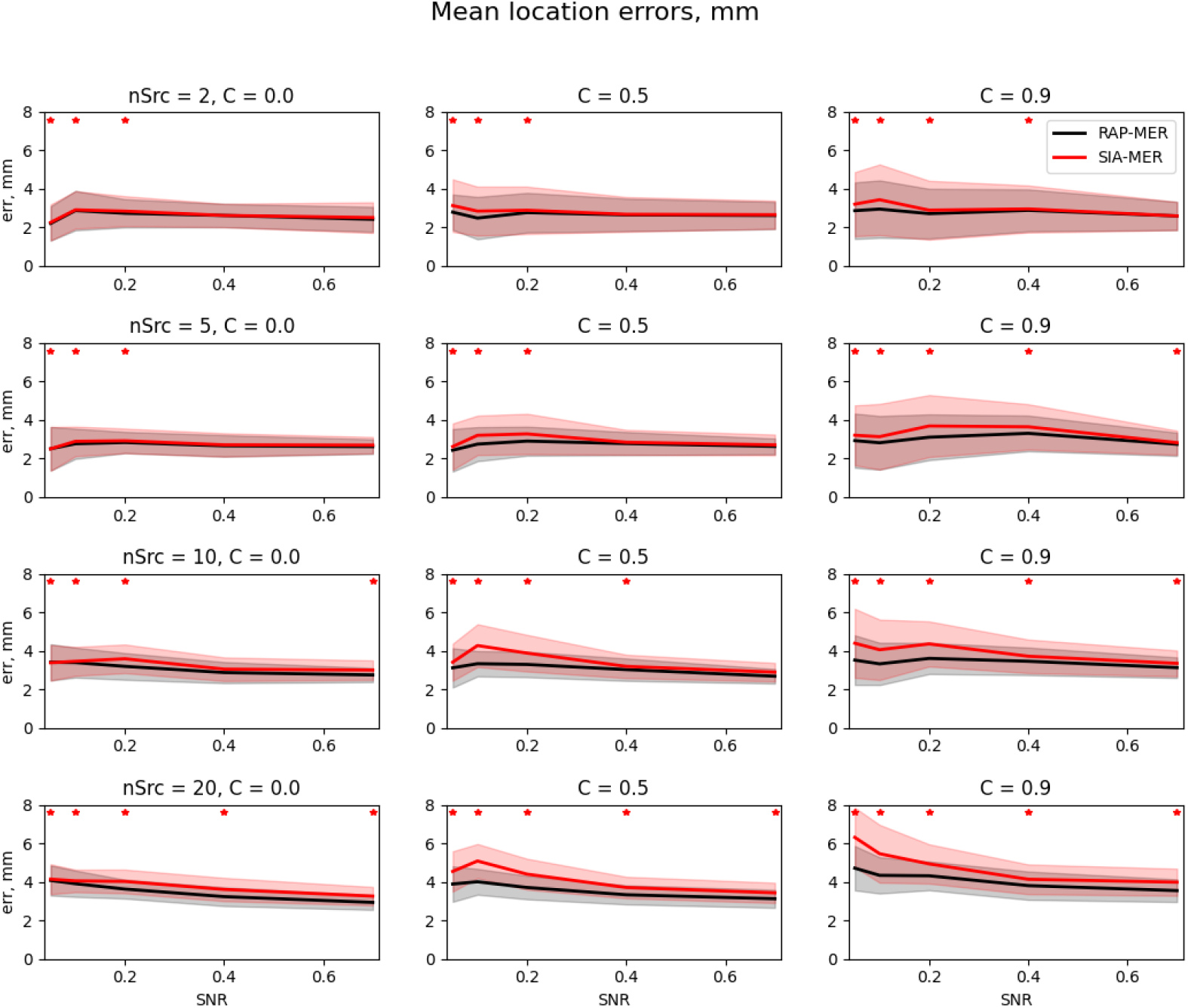
Average source localization errors for evoked (MER) **SMCMV RAP beamformer**(black) and **SMCMV SIA beamformer**(red) in the MEG pre-run noise case. The shadowed areas mark one standard deviation around corresponding curves. Asterisks denote SNR values where differences between the two methods are statistically significant.

**Figure S.8:**
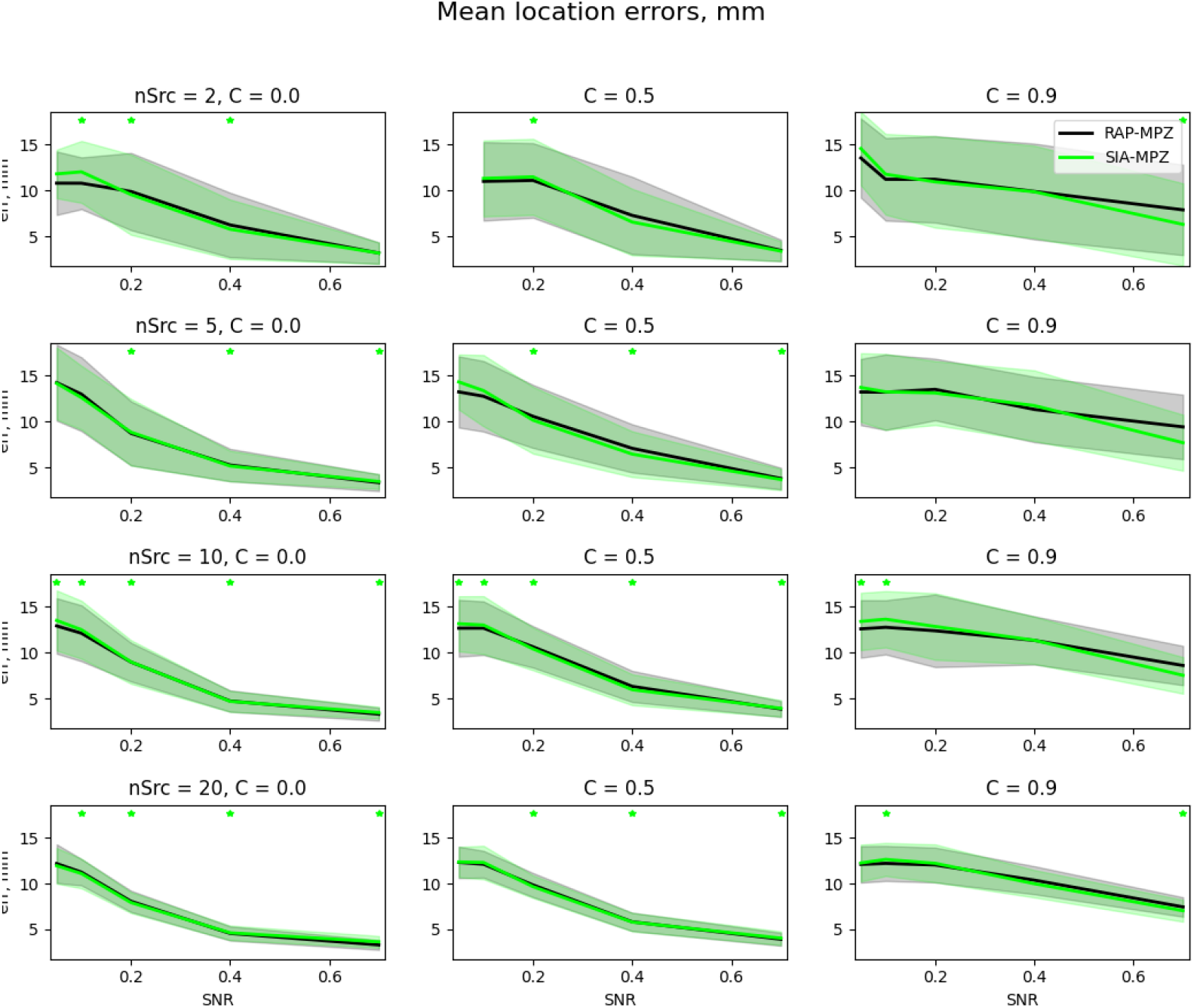
Average source localization errors for MPZ **SMCMV RAP beamformer**(black) and **SMCMV SIA beamformer**(green) in the MEG white noise case. The shadowed areas mark one standard deviation around corresponding curves. Asterisks denote SNR values where differences between the two methods are statistically significant.

**Figure S.9:**
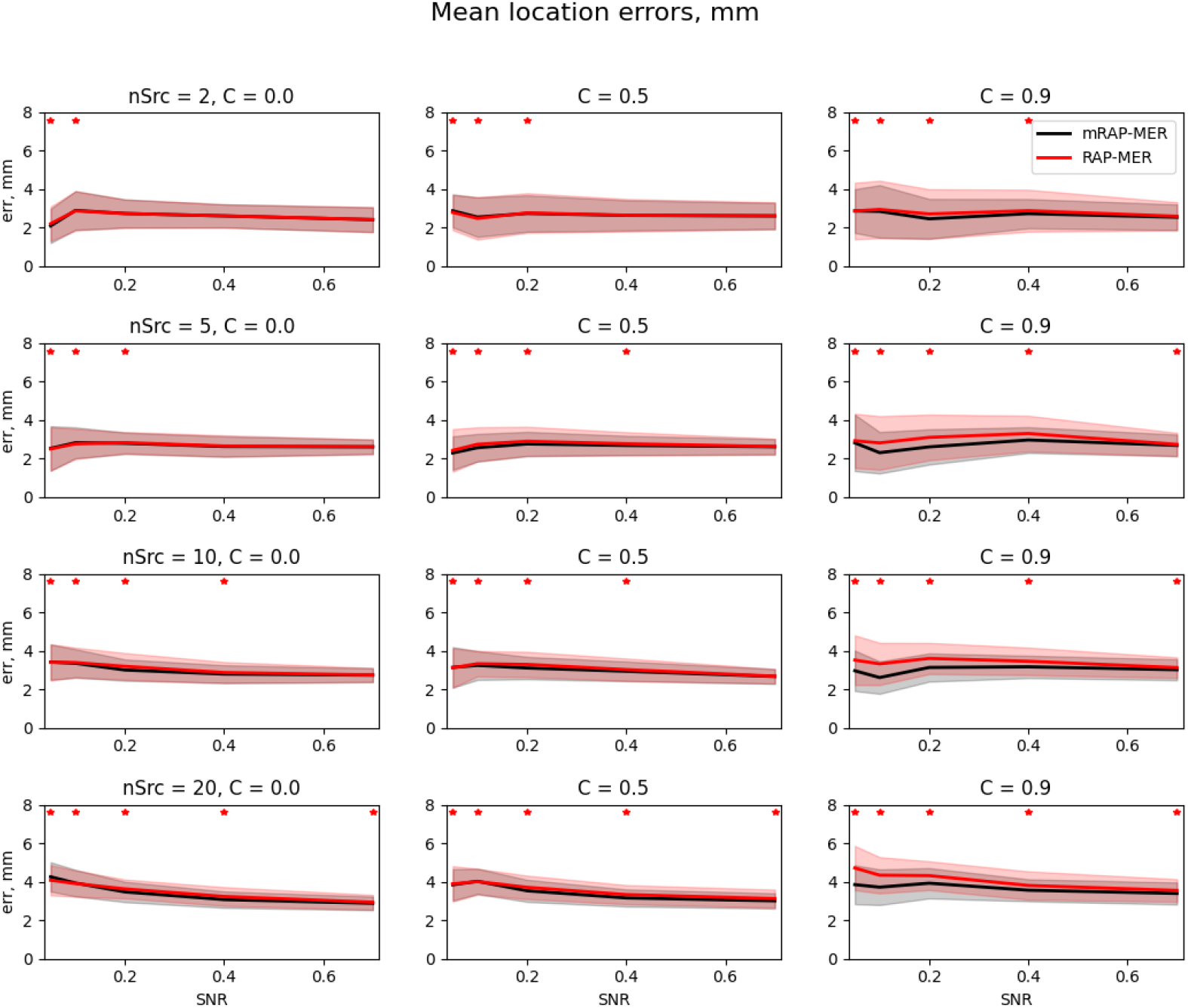
Average source localization errors for the MER **SMCMV RAP beamformer**(red) and **SMCMV modified RAP beamformer**(black) in the MEG pre-run noise case. The shadowed areas mark one standard deviation around corresponding curves. Asterisks denote SNR values where differences between the two methods are statistically significant.

